# Transmission of an environmental modification in quails across three generations: changes in the contribution to phenotypic variability

**DOI:** 10.1101/2025.07.04.663137

**Authors:** Stacy Rousse, Sophie Leroux, David Gourichon, Ludovic Calandreau, Andrea Rau, Sandrine Lagarrigue, Tatiana Zerjal, Frédérique Pitel, Sonia E. Eynard

## Abstract

**Background:** While epigenetic variations can contribute to shaping phenotypic diversity, it can be challenging to isolate and quantify the portion of trait variability under non-genetic influence. In this study we compared the phenotypic responses for different traits of two epilines of Japanese quails (*Coturnix japonica*) across three generations, using a large sample size. These epilines were built in parallel following (epi +), or not (epi −), an initial genistein ingestion in the ancestors’ diet and were maintained to harbour a similar genetic structure.

**Results:** Linear models were fitted to extract the fraction of variance allocated to multiple factors such as family, sex and epiline. The latter was found to be significantly associated with body weight. The contribution of the epiline to phenotypic variability progressively increased from the first generation (G0) to the last (G2), leading - for example in body weight at slaughter - to an average difference for adult males and females in G2 of 9 grams and 14 grams respectively, between epi + offspring and controls (epi−).

**Conclusions:** Although these findings suggest genetic drift, they could also reveal a possible transgenerational effect of the initial diet disruption. The analysis of other phenotypes displayed rare significant effects of the epiline. This innovative experimental design offered a unique opportunity to better understand the evolution of phenotypic variability and the parameters constituting it across three generations following or not an environmental change.

## Background

The increased frequency of environmental constraints due to climate change (e.g. heat waves, resource limitation …) can be a source of stress for animals. One of the key drivers of adaptation is the emergence of new phenotypes, which arise from the combined effects of genetic influence of parental alleles and non-genetic factors. Fluctuations in the parental environment can trigger non-genetic modifications, such as microbiota, learning or epigenetics, resulting in changes in gene regulatory mechanisms and thus gene expression, even in the progeny (David et al., 2019; Guerrero Bosagna et al., 2018). Understanding the impact and transmission of environmental effects to subsequent generations is crucial to comprehend how animals adapt to changing environments to maintain animal production needs while ensuring animal welfare. The study of a change in offspring phenotype, caused by an environmental signal experienced by the parental generation (and possibly previous ancestors), without involving genetic change in the offspring can be referred to as transgenerational plasticity in certain research fields (Holeski et al., 2012).

Most studies classically tackle the direct impact of the environment on the traits of exposed animals and on the performances of the subsequent generation. However, others have managed to evaluate a possible transgenerational non-genetic inheritance, considering an environmental effect to be transgenerationally transmitted when the individuals exhibiting trait variations were not exposed to the environmental stressor, even as germ cells (Skinner, 2008). On the contrary, an effect is considered as intergenerationally transmitted when trait variations are observed in the offspring of the individuals exposed to an environmental modification, therefore in an individual exposed as embryo or germ cell status. Multigenerational, including inter and transgenerational, studies in farm animals are challenging due to the need to raise multiple generations. To this end, Japanese quail (*Coturnix japonica*) is an ideal avian model for multigenerational analysis, thanks to its short generation interval, the wide range of measurable phenotypes, and the controlled breeding practices and limited space in which it can be reared. Previous studies on a variety of traits such as body weight, egg production and behavioural characteristics using Japanese quails as model organisms have reported multi and transgenerational phenotypic responses of the progeny after a thermal increase in the parental embryonic environment (Vitorino Carvalho et al., 2023), after in-ovo injection of genistein, an endocrine disruptor (Leroux et al., 2017), and after exposure to maternal stress (Charrier et al., 2024). These studies suggest a possible transmission of environmental effects through changes occurring in the prenatal embryonic environment. In addition, some evidence of the importance of grandmother’s diet was documented in mule and Muscovy ducks, with significant variations in ducklings and adult weights after methionine deprivation in the diets of the grandmothers (Brun et al., 2015). This suggests that diet disruptions can indeed be a cause of phenotypic response to an unknown yet memorised environment, through a specific inheritance mechanism.

In light of these previous analyses, we aimed to expand upon the pilot study and deepen our understanding of potential multigenerational effects. In the initial study (Leroux et al., 2017), the authors established two quail populations originating from a common founder line and differing only by an embryonic genistein exposure, using a mirrored single-pair mating design to minimize between-line genetic divergence. Genistein is a phytoestrogen compound naturally present in soy, and was notably associated with significant shifts in DNA methylation profiles in the promoter region of the Agouti gene in mice (Dolinoy et al., 2006). Among non-genetic factors influenced by the environment, DNA methylation, an epigenetic phenomenon, can be involved in the construction of phenotypes (Duncan et al., 2014). Several studies in mice and humans have demonstrated the influence of genistein on DNA methylation, including in embryonic stem cells (e.g. Day et al, 2002, Sato et al 2011, Bilir et al, 2017). Genistein is also known, among others isoflavones, to affect performances in livestock, including poultry (see Grgic et al, 2021).

As genistein has been shown to induce stable DNA methylation changes and to affect several traits in different species, it was considered as a relevant chemical compound to study environmentally induced epigenetic variation in a livestock species. The pilot experiment showed that the embryonic environment can affect growth, reproductive and behavioural phenotypes three generations later and was associated with changes in blood DNA methylation (Cerutti et al., 2023), thereby suggesting a contribution of non-genetic inheritance mechanisms. The two populations, issued from treated or control ancestors, were named ‘epilines’ to emphasise the potential epigenetic inheritance of environmental influences during development, even if such putative non-genetic transmission may be due to other non-genetic processes such as cytoplasmic, microbiotal or behavioral transmission (Bonduriansky and Day, 2009; Day and Bonduriansky, 2011, David et al 2019,). However, in this pilot experiment, genistein was directly injected into the egg and intermediate generations were not phenotyped or characterised at the molecular level, which limits inference to realistic field-like conditions and prevents a fine dissection of multigenerational responses.

In addition, the use of genistein as supplementation within the feed of Japanese quails has been shown to lead to better adaptation of birds to heat stress by stimulating feed efficiency and growth rate (Onderci et al., 2004). We therefore designed a multigenerational quail experiment in which maternal dietary genistein supplementation acts as the environmental exposure. This design aims to more closely mimic living conditions (ingestion rather than injection), while generating populations suitable for joint genetic and non-genetic analyses across all generations up to F3. We hypothesised that maternal genistein supplementation would induce specific phenotypic and epigenetic changes in directly exposed offspring, and that part of these changes would persist in unexposed descendants, consistent with non-genetic inheritance of environmentally induced variation.

While epigenetic variations can contribute to shaping phenotypic diversity in offspring, isolating and quantifying the portion of trait variability influenced by epigenetics remains challenging. To date, although genetic contribution has largely been documented for a wide range of traits in a large number of poultry species, no direct quantification of the epigenetic influence on these traits has been reported. The environment usually accounts for a large part of the observed variability depending on the phenotypes measured, which translates to low heritabilities even under selection (Chomchuen et al., 2022; Gaya et al., 2006; Lormant et al., 2020; Prince et al., 2020; van Oers et al., 2004; Wright and Henriksen, 2020) providing a compelling reason to estimate the contribution of non-genetic factors, including epigenetics, to trait variability. In this study, a case - control design allowed building two distinct epilines. The epiline information provides an indirect way to estimate the epigenetic contribution to the trait, as well as other non-genetic factors. By investigating the multigenerational transmission of the effects of an initial diet disruption across three generations of quail, we aimed to capture the inter- and transgenerational impacts of environmental influences, likely mediated by non-genetic mechanisms. The portion of phenotypic variability linked to an ancestor’s environmental change and its evolution across generations was estimated in body weights.

## Methods

### Quail mating design

All animals in the experiment come from a Japanese quail (*Coturnix japonica*) experimental line (HSR, for High Social Reinstatement) arising from a divergent selection on social motivation (Mills and Faure, 1991). From this common genetic background, twenty initial founding families were crossed, producing the generation G-1. In each founding family, two full sib females were mated at maturity with the same male, originating from another family. One of the females was administered a genistein capsule (100 mg, Molekula group) for thirty days after mating while the other one was only provided with empty capsules, leading to the development of G0 individuals in two different embryonic environments. The females that received the genistein are the dams of the treated epiline (thereafter called epi+), in opposition to the control epiline (epi-). After this initial diet disruption, the quails were mated following a mirror design, with males and females from the same founding families mated in each line, to maximize the genetic balance between the two epilines. The goal of this mating design is to maintain the quails in both groups as genetically similar as possible while ensuring that the only potential difference between them is the initial dietary disruption. Therefore, if phenotypic variability is observed between the two groups, it can be hypothesized that it results from the change in feed intake during the first generation and limits the possibility that it arises from a genetic factor. Groups were then maintained under standard rearing conditions, at the breeding facility, for the three subsequent generations (G0, G1 and G2), with water and feed ad libitum (Fig.1). The ‘epiline’ terminology is therefore used to refer to these two parallel groups of quails whose distinction lies within their non-genetic factors, that includes epigenetic mechanisms *per se* as well as any other environmental effects that can be transmitted across generations. From hatching to 49 days-of-age, individuals were floor-raised on litter altogether, in climate-controlled housing. Animals were then placed within cages of two individuals of same sex, until the age of twenty weeks, when they were coupled for mating. All diet, temperature and light conditions were standard and the same for all animals, all reared in the same barn.

**Figure 1.**
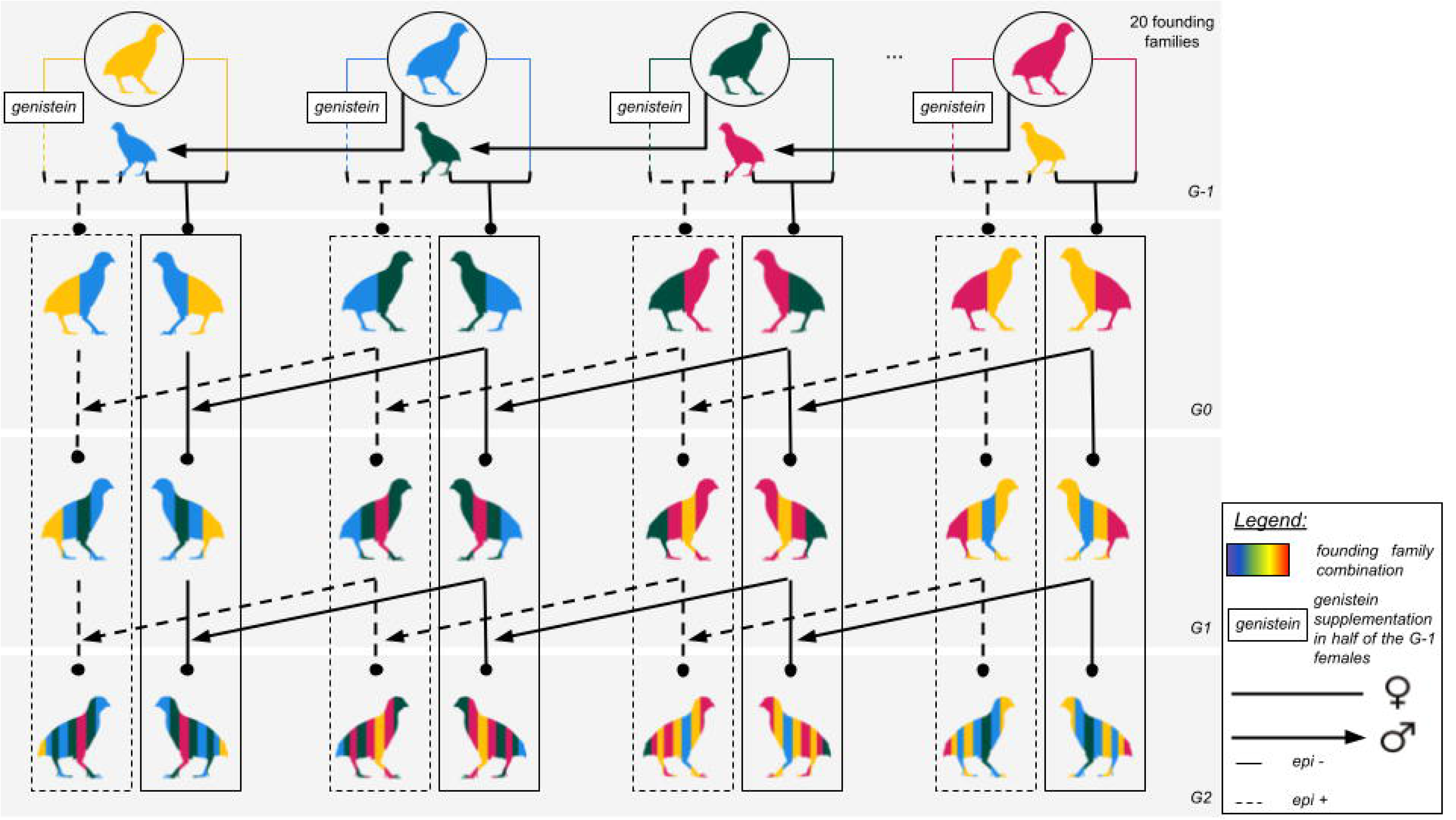
Quail mirror mating design. The quail mating design runs over four generations of quails, the first one (G-1) containing the founding populations that have been split into two populations from G0 onwards and mated independently for three successive generations. Genistein capsules were administered to females in G-1 while their full-sib female counterparts were provided empty capsules. Both dams were then mated with the same male coming from another genetic background. The G0 progeny was divided in 2 epilines (epi+ deriving from the dams having ingested the genistein and epi-deriving from the control dams) of similar genetic background. The following matings were controlled to respect the balance of the genetic contribution from founders by ensuring a mirror between epilines (solid lines: epi- and dashed lines: epi+). The theoretical genetic contribution of founders is represented by the colors.

### Phenotypic traits

Phenotypic records were collected over four generations for body weights, tissue weights, egg production and behavioural response to isolation.

#### Body weights (BW)

Body weights were measured at different developmental stages: one week, four weeks, seven weeks and slaughter (thirty weeks of age on average) for all quails. A unique scale was used for all measurements and weights were collected at the same age when possible to ensure consistency between generations. Slaughter was done through electronarcosis immediately following the recording of the final body weight measure.

#### Tissue weights (TW)

Several tissues were weighed: abdominal fat (for all generations), heart, liver (for G0 onwards), and testicle (for G1 onwards). All weights were adjusted by correcting for the weight of the quail at slaughter.

#### Egg production (PROD)

At twenty weeks of age, quails were placed in cages for reproduction. Egg production data, total egg number during the laying period and precocity, were recorded from the first day in the cage for 49 days. The laying rate was estimated as the number of eggs laid over the laying period. The mean egg weight per quail was calculated on a basis of four eggs collected and weighed per quail.

#### Behaviour at isolation (BHV)

Adult quails underwent an open field behavioural test in which their reactivity to a new environment was assessed. Each quail was placed in a cubic arena (80 cm) of white wood with a floor made of beige linoleum, in an unknown experimental room surrounded by an unknown white curtain. Each quail was individually placed at the centre of the arena and allowed to freely explore it for 300 seconds. For each trial, the time spent in the inner area (a square of 20 cm × 20 cm at the centre of the arena), the frequency of time spent at the periphery (outside the inner area) as well as the total distance covered by the quail was recorded by a camera placed above the arena. The open field test was performed for all generations.

### Statistical analysis strategy

The phenotypes were analysed following these steps: first, we performed a descriptive overview of the phenotypes by looking at the overall trait distribution for all raw phenotypes, the distributions per generation and sex within each epiline are shown in Supplementary file 6; second we tested normality; third we looked at the potential impact of the epiline factor by mean comparison; fourth, we applied some transformations on our phenotypes of interest when necessary; and at last we ran a linear model to obtain an estimation of the part of variability explained by different factors of interest.

#### Normality and mean comparison tests

Shapiro-Wilk normality tests were performed on all raw phenotypes. Since all variables did not meet the normality assumptions, Kruskal-Wallis tests were applied to perform pairwise mean comparisons between epilines. Given the strong sexual dimorphism in quails, the test was applied to all raw phenotypes in each sex and generation separately. The null hypothesis of H0 “No significant effect of the epiline on the phenotype” was rejected at the 5% level of significance. The results are shown in Supplementary file 6.

#### Estimation of the contribution of parameters to body weight traits

Following the results of Kruskal-Wallis tests, we decided to focus our analyses on body weight traits (results on other traits are shown in Supplementary file 7). To estimate the percentage of variability explained by influential factors such as the family, sex and epiline, linear models were fitted to normalized body weights.

#### Data normalization

To avoid biased estimates of coefficients and meet the assumptions of linear modeling, we calculated the skewness of all traits following different transformations using the R ‘moments’ package (version 0.14.1). Skewness of Log10, square root and inverse transformed data were compared to that of the raw data and the transformation associated with the lowest skewness value was subsequently retained for each trait (Supplementary file 6). The choice of transformations tested were selected among commonly applied data transformations prior to linear modeling (Fink, 2009).

#### Linear models (LM)

For each body weight at each generation, the optimal set of parameters was first determined with a stepwise model selection using the stepAIC function of the R ‘MASS’ package (version 7.3-65). A linear model was fitted for each trait as follows:

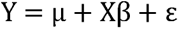

where Y is the phenotype, µ is the overall mean, X is the design matrix for the fixed effects, β is the coefficient vector of fixed effects and ε is the residual. Fixed effects included in the model were sex (male, female), epiline (epi-, epi+) and family (origin from founders) and their pairwise interactions (sex:epiline, sex:family and epiline:family). Effects were iteratively added or deleted until the model with the lowest Akaike information criterion (AIC) was identified for each trait. The LM fit on the other traits is shown in Supplementary file 7.

#### Cross Validations

After selecting the optimal model, we evaluated the fit of the model using 100 random fold cross validations (80% of data in the Train set, 20% in the Test set). For each fold, the Train set was used to fit the model using the optimal equation determined with stepAIC, and it was then used to predict the phenotypic response of the Test set. Correlations between observed and predicted Test phenotypes were then assessed. The coefficient of determination R², which represents the overall variance of the trait captured by the model, was computed for each fold.

#### Effect of parameters

The proportion of phenotypic variability explained by each factor for a given trait was estimated using an analysis of variance (ANOVA – type III). The total sum of squares SS_Total_ is decomposed into the sum of squares for each factor j SS_Factor_j_ and residual variance SS_res_. The proportion of variance explained (PVE) of factor j may be calculated as:

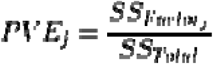

Estimates were obtained using the R package ‘FactoMineR’ (version 2.12) with Type III sum of squares. We ensured success in variance components estimations by retaining only families with a sufficient number of quails per epiline.

All statistical analyses were performed using R 4.5.2 (R core team).

## Results

The summary statistics of all traits collected from all generations are available in Supplementary Table S1.

### Impact of the epiline on phenotypes and evolution across generations

#### Body weights (BW)

Significant differences between epilines were observed across several body weights. Females showed significant differences starting from generation G1 at all ages with epi+ quails systematically weighing more on average than controls epi-. The same pattern was found in males where the groups began to consistently diverge in G2 at all ages; however, no significant difference was observed in earlier generations (Fig. 2).

**Figure 2.**
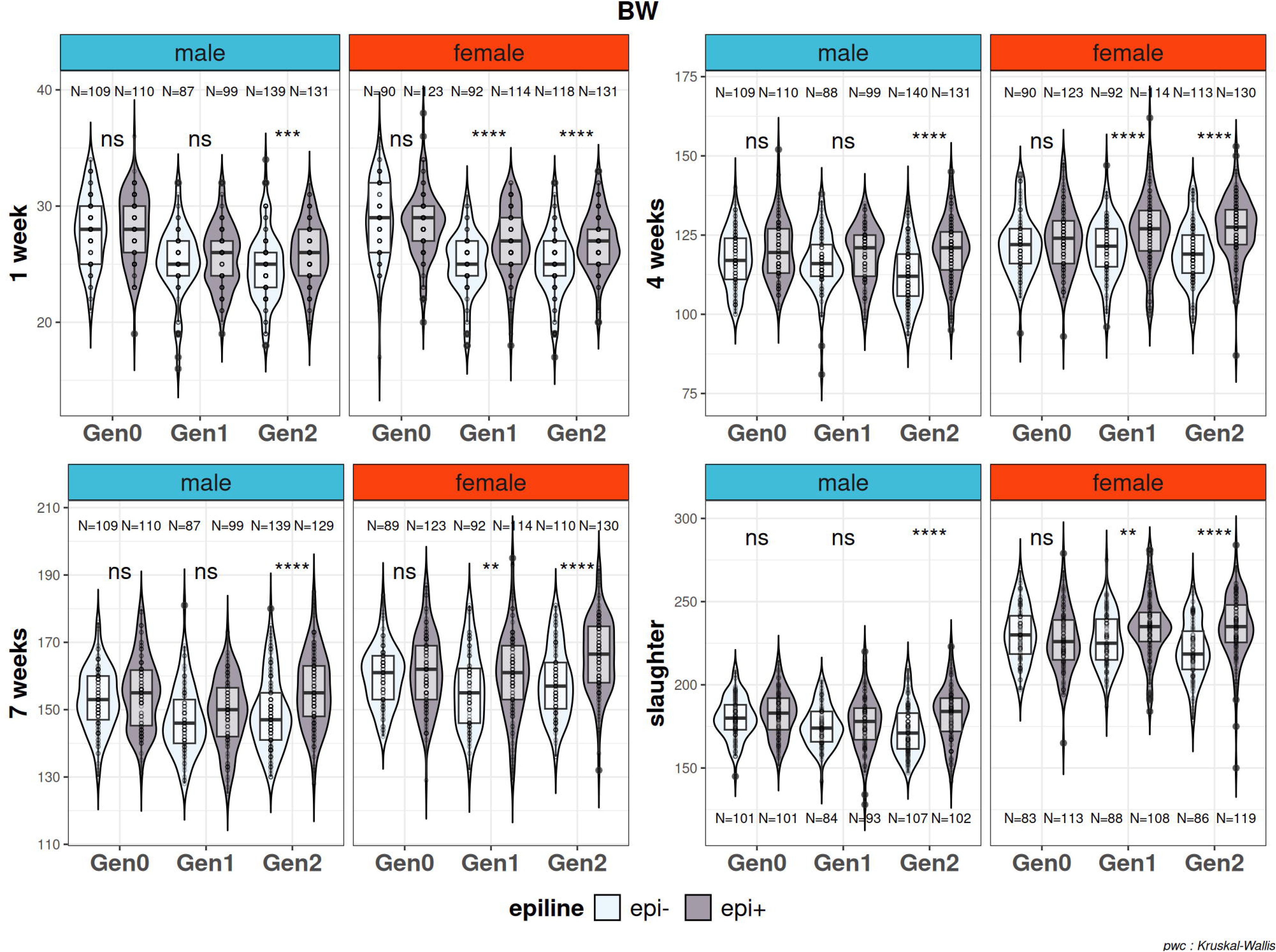
Mean comparison of body weights at one week, four weeks, seven weeks and at slaughter (thirty weeks) between epilines. Kruskal-Wallis tests were performed to identify differences between epilines for each phenotype, on raw estimates, for each sex and generation independently. N values indicate sample sizes within each sex and epiline per generation. The asterisk indicates the level of significance of the Kruskal-Wallis test results, ns p-value > 0.05, * p-value < 0.05, ** p-value < 0.01, *** p-value < 0.001 and **** p-value < 1e^-4^.

Differences in average BW at all ages became increasingly more pronounced over the generations. In G0, the difference between epi+ and epi- at slaughter was on average 4 grams for both sexes, which was not found to be significant. In G1, the difference between epilines for females was on average 7 grams and reached 14 grams in G2. In males, a similar pattern was observed in G2, with epi+ quails weighing 9 grams more on average at slaughter whereas a maximum difference of 2 grams was observed in earlier generations. The growth curve to highlight this longitudinal evolution is added in Supplementary Figure S1.

#### Other phenotypes

For other phenotypes, such as tissue weights (TW), egg production (PROD) or behaviour at isolation (BHV), significant differences between epilines were infrequent and sometimes linked to individual singularities (adjusted heart weight in G2 males, frequency at periphery in G2 males, precocity in G1 females). Adjusted heart weight in G1 males, duration spent at the centre of the arena in G1 males, laying rate in G2 and precocity in G1 were the only traits for which a significant difference between epilines was detected. These results are displayed in Supplementary Figure S2-S4.

### Body weight variability explained by parameters: evolution across generations

#### Effect of the epiline

Model fitting revealed that the epiline explained a different portion of the variability observed in BW according to the generation and age of the quails. The epiline effect was found significant from four-weeks old quails onwards, in all generations. First, at one week of age, results showed significance in G2 only, with the epiline explaining 7.6% of the variability. Then, for four- and seven-weeks old animals, the epiline explained a negligible amount of weight variability in G0 (under 1%), but it was found to increase in later generations, accounting for around 2% in G1, to finally reach a more substantial proportion in G2 (over 15%) at those ages. Finally, the contribution of the epiline to BW in adults (BW at slaughter) was much lower than at previous ages (less than 1% in G0 and G1 and 4.5% in G2) (Table 1, Fig.3).

**Figure 3.**
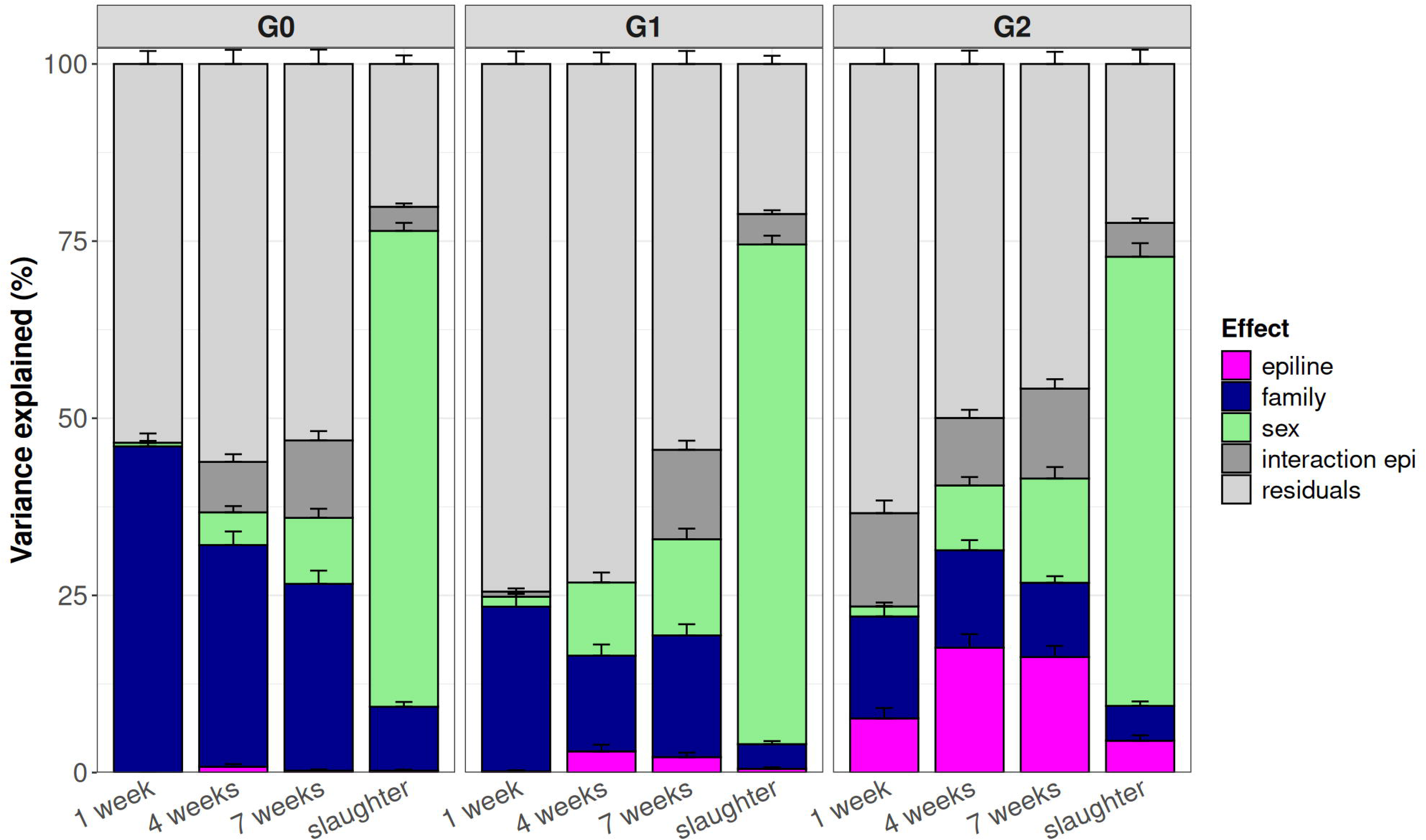
Contribution of parameters (sex, family, epiline, and interaction) to body weight variability (in percentage) per generation. Portion of phenotypic variability explained by the effect of sex, family, epiline and their interaction as extracted from the best linear model for body weight traits. The interaction parameter always involved the epiline (hence the terminology ‘interaction epi’ used). More specifically, it refers to: epiline:family for BW at 1 week (G2), BW at 4 weeks (G0, G2), BW at 7 weeks (G0, G1, G2) and BW at slaughter (G1, G2); sex:epiline for BW at 1 week (G1); sex:epiline and epiline:family for BW at slaughter (G0). Body weights throughout growth are arranged within generations G0, G1 and G2.

**Table 1.** Estimates of the contribution of parameters (and standard deviation) to body weight variability (in percentage).

| Trait | Generation | epiline | family | sex | interaction | residual |
| --- | --- | --- | --- | --- | --- | --- |
| 1 week | G0 | 0 (0) | 46.037 | 0.515 | 0 (0) | 53.448 |
|  |  |  | (1.819) | (0.257) |  | (1.829) |
|  | G1 | 0.153 | 23.264 | 1.396 | 0.732 | 74.454 |
|  |  | (0.176) | (1.794) | (0.512) | (0.441) | (1.777) |
|  | G2 | 7.641 | 14.389 | 1.414 | 13.17 | 63.386 |
|  |  | (1.467) | (1.449) | (0.552) | (1.783) | (2.279) |
| 4 weeks | G0 | 0.818 | 31.288 | 4.617 | 7.119 | 56.158 |
|  |  | (0.376) | (1.912) | (0.881) | (1.084) | (1.990) |
|  | G1 | 2.984 | 13.513 | 10.322 | 0 (0) | 73.181 |
|  |  | (0.946) | (1.573) | (1.410) |  | (1.634) |
|  | G2 | 17.607 | 13.761 | 9.146 | 9.526 | 49.961 |
|  |  | (1.923) | (1.432) | (1.190) | (1.151) | (1.885) |
| 7 weeks | G0 | 0.264 | 26.363 | 9.315 | 10.927 | 53.131 |
|  |  | (0.183) | (1.873) | (1.284) | (1.315) | (2.027) |
|  | G1 | 2.175 | 17.177 | 13.574 | 12.620 | 54.454 |
|  |  | (0.625) | (1.575) | (1.500) | (1.291) | (1.837) |
|  | G2 | 16.304 | 10.482 | 14.708 | 12.691 | 45.815 |
|  |  | (1.568) | (0.920) | (1.621) | (1.327) | (1.713) |
| slaughter | G0 | 0.262 | 9.026 | 67.148 | 3.411 | 20.153 |
|  |  | (0.145) | (0.677) | (1.141) | (0.466) | (1.204) |
|  | G1 | 0.530 | 3.479 | 70.513 | 4.308 | 21.170 |
|  |  | (0.209) | (0.429) | (1.247) | (0.521) | (1.153) |
|  | G2 | 4.492 | 4.917 | 63.378 | 4.789 | 22.423 |
|  |  | (0.758) | (0.628) | (1.917) | (0.618) | (2.010) |

#### Effect of other factors

Contrary to the increase of the epiline effect across generations, the portion of phenotypic variance explained by the family declined, starting, at one week of age, from 46% in G0 to 23% in G1 and decreasing again to 14% in G2 (Table 1).

The family effect was also stronger in younger animals. It ranged from 46% to 9%, from 23% to 3.5%, and from 14.4% to 5% between one-week of age and slaughter for G0, G1 and G2 respectively. On the other hand, the proportion of variability explained by the sex always increased with animal age. It ranged from 0.5% to 67%, from 1.4% to 70.5% and from 1.4% to 63.4% between one-week of age and slaughter for G0, G1 and G2 respectively. The interaction parameter, when conserved, always involved the epiline and the family, except in two traits: in body weight at one week in G1 where it concerned epiline and sex, and in body weight at slaughter in G0 where it concerned epiline and sex as well as epiline and family. Depending on the generation and the trait, this parameter showed little to moderate contribution to phenotypic variability, ranging from 0 to 12%. Overall, the residual remained high in all traits but BW at slaughter, regardless of the generation.

#### Cross validations

The predictive performance of the models, evaluated using cross validations, varied considerably across BW traits. Body weight at slaughter consistently exhibited the highest prediction accuracy across all generations (R² ≥ 0.8, Corr ≥ 0.8), highlighting good model performance for adult individuals. For younger animals, the coefficients of determination were significantly lower than at slaughter despite remaining of reasonable magnitude (R² ≤ 0.5 in G0 and G1 and R² ≤ 0.6 in G2). The correlations between Train and Test sets were moderate, ranging between 0.4 and 0.6, suggesting limited predictive power of our models for these traits; these results are in line with the large residual variance estimated (Fig. 4).

**Figure 4.**
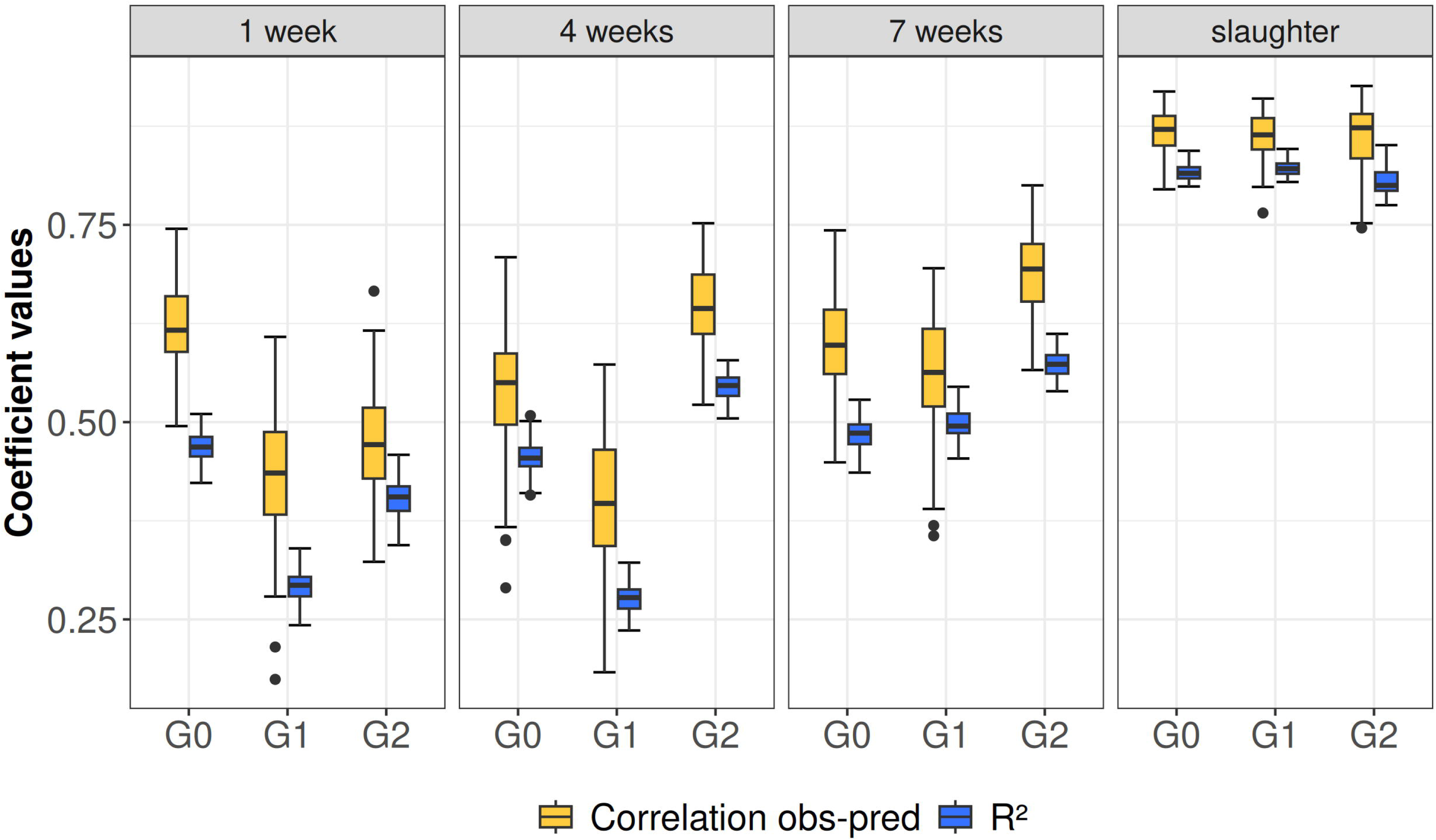
Model robustness (R²) and correlations after cross validations. The coefficient of determination R² (blue) and the correlation between observed and predicted values in the Test set (yellow) were determined in 100 cross validations, to evaluate the predictive performance of linear models. The results show the distribution of R² and correlation values in each of the 100 cross validations, and are displayed per generation (x-axis) for each body weight (1 week, 4 weeks, 7 weeks and at slaughter).

#### Other phenotypes

Results on the estimation of the effects of parameters on the variability of other traits is shown in Supplementary file 7. The epiline was a significant factor to keep for explaining a portion of phenotypic variability for several traits. In adjusted tissue weights like heart, abdominal fat and testicle, this parameter was kept after the stepAIC in all generations. For the liver, it was only found significant in G2. For production traits, the epiline was found significant in precocity and in adjusted egg weight at every generation but not for the laying rate. Finally, in behavioural traits, the epiline factor was kept in the models for total distance and frequency at periphery in G1 and G2 only, whereas it was kept in G0 and G1 for duration at centre. Although the epiline appears as a significant factor in the aforementioned traits, the estimates of the portion of variability explained remain below 5% in all, and negligible in most (inferior to 1% in adjusted abdominal fat, adjusted liver weight, total distance, frequency at periphery and duration at centre). Besides, for all the traits presented in Supplementary file 7 (D), a substantial part of phenotypic variance remains uncaptured by our model, highlighting a strong limit in the explanatory power of our models and a limited confidence towards the estimates of the portion of phenotypic variability explained by the epiline for these traits.

## Discussion

The aim of this study was to analyze phenotypic variations across three generations of Japanese quails (*Coturnix japonica*) following supplementation with genistein, an endocrine disruptor. For this purpose we built one of the largest available designs in livestock containing groups impacted (treatment) or not (control) by environmental stressors and multiple generations of mirror breeding to limit the impact of genetic variation in the determination of trait changes through time.

### Genistein ingestion is associated with different phenotypic responses across generations

In both sexes, body weight (BW) tended to increase in the treated epiline (epi+) compared to the control epiline (epi-), with the difference between epilines becoming progressively more pronounced from generation G0 to G2. In a pilot study (Leroux et al., 2017), a similar pattern was observed in G2 females: genistein injection into the egg was associated with increased BW at three weeks of age. However, in that study, the effect was reversed in males (genistein injection associated with decreased BW). In the present study, we did not observe such sex-specific interactions regardless of the generation or age considered. On the other hand, the results obtained here concur with other studies regarding the significant impact of genistein on BW. The scientific community remains divided on the impact of the phytoestrogen on the organism (Rockwood et al., 2019; Xiang et al., 2023), with most studies tackling the direct effect of ingestion on the concerned individuals. More generally, body weight management is of importance in poultry, as this trait often correlates with production and reproductive efficiency (Robinson et al., 1993). The choice of feed can be critical to optimize carcass related traits in broilers offspring (Mahmoud Ali and Hamid Hassan, 2022) as well as ensuring ideal reproductive performances (Aldahmani and Rasah, 2026). In that regard, our results on body weight divergence provide insight on the need to vigilantly monitor feed intake as well as feed source, according to the breeding strategies at stake.

Production (PROD) traits did not exhibit the strong epiline-related effects seen previously in the pilot study, where epi+ females began laying eggs more than eight days later than control females on average. Likewise, abdominal fat weight and behaviour (BHV) traits did not demonstrate the previously reported significant differences between epilines. The discrepancies between the two studies, despite using the same quail genetic line and the same endocrine disruptor, are likely attributable to differences in the method of genistein administration: in the pilot study, 500 µg of genistein were directly injected into the egg before incubation, whereas in the present study genistein was administered orally to the G-1 mother to mimic more realistic breeding conditions, resulting in substantially lower concentrations of the molecule in the egg (Lin et al., 2004).

Direct effects of genistein ingestion on phenotypes have been reported in humans and animal models. Although this molecule offers promising results in cancer treatment (Casari et al., 2025; Rasheed et al., 2022), soy overconsumption has demonstrated harmful effects on reproductive (Liu et al., 2020; Lozi et al., 2021; Rashid et al., 2022; Toktay et al., 2020) or cognitive (Svensson et al., 2023) functions. These effects of endocrine disruptors on health seem to be persistent over several generations, although this remains relatively unexplored, despite a few specific studies (Brulport et al., 2021). The molecular effects of this phytoestrogen have been characterized, especially in cancer research, genistein acting similarly to estrogens, but also as an antioxidant, anti-inflammatory agent and inhibitor of cell cycle regulators (see Naponelli et al., 2025; Sharifi-Rad et al., 2021 for a review). These studies also underline the low bioavailability of the molecule, especially in its glycosylated form, as reported for other flavonoids (Ross and Kasum, 2002). A similar issue arises in quail, where it was observed that glycosylation reduces the transfer and accumulation of genistein in egg yolks after maternal supplementation (Lin et al., 2004), indicating that the chemical form and dose influence embryonic exposure. Despite these pharmacokinetic constraints, it has long been known that supplementing the diet of broilers with soy isoflavones, including genistein, improves growth performance, especially at low doses (< 80 mg/kg) (Kamboh and Zhu, 2013; Rasouli and Jahanian, 2015).

Maternal dietary supplementation with genistein has also been shown to affect offspring growth: in broilers, low maternal doses (e.g. 40 mg/kg) increase embryo weight, while higher doses (e.g. 400 mg/kg) can decrease it, indicating a nonLJlinear, doseLJdependent response (Lv et al., 2018). In quail, our previous work using direct egg injection of genistein, which corresponds to a high dose in the embryonic environment (500 µg), showed reduced body mass at several ages, confirming that high developmental exposure can alter growth trajectories (Cerutti et al., 2022). The divergence in body weight trend observed between the pilot study (Leroux et al. 2017) and our study confirm the dose effect on this trait. In fact, the injection practiced in Leroux et al. 2017 led to a reduction in offspring weight for the treated group whereas the ingestion, this study, shows an increase. Adding genistein to the maternal diet has been shown to increase genistein concentration in egg yolks, in Japanese quail, confirming that the molecule may reach the embryo through the yolk (Akdemir and Sahin, 2009; Lin et al., 2004). Our oral maternal route thus ensures realistic embryonic transfer via yolk, supporting phenotypic effects while allowing tests of non-genetic inheritance—our core question.

Overall, results reveal different sensitivities of the phenotypes towards genistein ingestion. Although this molecule is already used as a supplement to increase BW in broilers (Lv et al., 2018), it has been observed that it affects other phenotypes as well, but not as strongly (Navarro-Guillén et al., 2025).

### Missing variability highlights the complexity of phenotypes

The proportion of phenotypic variance explained by the family displayed a clear trend with age, decreasing over the lifetime (with the exception of week 7 in G1). In our model, the family factor was accounted for as a way to include the ancestral genetic effect. This suggests that the influence of the inherited genetic background becomes progressively less determinant as individuals age. Conversely, the contribution of the sex factor increased with age, consistent with the emergence of pronounced sexual dimorphism in quail. These patterns illustrate that different factors influence animal phenotypes in distinct ways at different stages of life, and that the proportion of phenotypic variance explained by a given factor is therefore strongly dependent on the individual’s age at the time of measurement.

Nevertheless, residual variance was substantial across all body weights and this is even more valid for other traits. This is highlighted by the generally moderate values of coefficient of determination (R²), suggesting the difficulty of the models to accurately reflect the overall variability of BW, especially in early life. The R² values for body weight at one week (< 0.5 in all generations) suggest limited explanatory power of the model. Given the low effects of the epiline estimated for this trait, the reliability of the estimation is questionable. In poultry, a major effect of the dam is expected to influence most early-life traits of the offspring through egg size and composition (Halbersleben and Mussekl, 1922; Widowski et al., 2022). In our model, this dam effect is absorbed within the family factor, and our results reveal that it is the factor which contributes most to body weight variability at one week of age, for all generations. So, in spite of its limited explanatory power, the model seems to show coherent trends with biological meaning. Over time, especially after four weeks of age, R² increases slightly showing a better suitability of our model. A potential explanation for the large residual variance relies on the complex determinism of phenotypic traits collected, influenced by various environmental factors, some of which the design may have failed to eliminate despite the controlled management. Multiple studies also reported high residual variances remaining, for example on a New Zealand bird species, a study found that home-range sharing and genetic relatedness among individuals contributed to phenotypic similarities for several morphological traits, yet residual variance remained high (Rutschmann et al., 2020). Similarly, Béziers et al., 2019 reported high residual variance for stress-induced corticosterone levels in a barn owl population, hypothesizing an additional factor beyond genetics or nest environment: the experienced environment. Since all individuals from the same generation in our study were reared under identical conditions, a plausible explanation for this residual variance would be the transmission of the experienced environment from the first generation, possibly involving slight variations in genistein dosage among eggs. The embryonic environment is known to influence the individual phenotype later in life, especially in birds (Frésard et al., 2013), and our results, even with a limited explanatory power of the model, tend to confirm that non-genetic factors contribute to a wide range of traits.

### Possible inter and transgenerational transmission of genistein effect

The availability of phenotypic records for all individuals of the design highlighted an overall progressive increase of the epiline effect from G0 to G2 on BW. This phenomenon may be due to genetic drift (Falconer and Mackay, 1996; Hospital and Chevalet, 1996), even though the “mirror” mating design was implemented to minimize genetic effects, ensuring a balanced origin of founders between epilines in each generation. David & Ricard 2023 showed a potential impact of drift on the phenotypic difference observed between epilines after three generations on a simulated design similar to that used in our experiment. By running additional simulations across all generations of interest (Supplementary file 8) we show that the phenotypic difference observed between epilines (from G1 onwards for females and in G2 for males) in our experimental design appears greater than when simulating drift.

Alternatively, one could explain such change through an increasing environmental influence across generations. Recently, the impact of the parental epigenome on offspring’s developmental phenotypes has been a research topic of growing interest in different livestock species like cattle (Kiefer et al., 2021), pig (Wu et al., 2024), sheep (Fonseca et al., 2023) and poultry (Hu et al., 2020). One might expect non-genetic (particularly epigenetic) differences between epilines to decrease progressively due to epigenetic reprogramming in the germline (Burton and Greer, 2022). This resetting is not so clear in chicken regarding DNA methylation (Kress et al., 2024). This may partly explain the observed multigenerational effect, as partial reprogramming may allow some epigenetic signatures to escape and be transmitted across generations (Heard and Martienssen, 2014).

Surprisingly, epigenetic effects can even strengthen over generations despite germline resetting. For example, ancestral vinclozolin exposure in rats seems to induce sperm methylation changes that persist beyond 13 generations, correlating with rising disease incidence (Korolenko et al., 2025). Repeated exposures further amplify multigenerational adaptations, as seen in *C. elegans* (Burton et al., 2020; Burton and Greer, 2022, see Liberman et al., 2019) or already reported in quail (Vitorino Carvalho et al., 2023). Even without repeated challenges, effects may intensify via secondary genetic modifications as observed in rats treated with vinclozolin (Skinner et al., 2015).

More generally, transgenerational studies conducted on different models reveal that environmental exposures leave epigenetic marks that are partially reset in the germline, but can yield progressive phenotypic changes through genetic or non-genetic mechanisms (Burton and Greer, 2022). Within this framework, our observations may reflect gradual amplification of initial non-genetic perturbations in specific genomic or epigenomic regions, leading to an increase in the proportion of variance explained by epiline across generations, even post-exposure. Given the unclear transmission mechanism of environmental effects in birds, monitoring epigenetic changes throughout generations using molecular data would shed light on the molecular dynamics at play to verify this assumption in Japanese quails (*Coturnix japonica*).

Indeed, in this study, genistein was only administered once and individuals were not selected, suggesting a possible transmission of the environmental effects to the progeny. This hypothesis concurs with results found by Karami et al., 2025 on the same dataset. Indeed, the authors designed an additional covariance for offspring descending from either epi+ or epi-individuals, as they share an extra similarity due to their embryonic environment. The inclusion of this additional covariance was significantly different from 0 for BW traits,, marking a probable transmission of genistein effects to the progeny. Their result supports a potential transmission of environmental effects via non-genetic mechanisms.

Indeed, several experimental studies show that genistein can modulate the epigenome, which offers a plausible mechanistic link between early exposure and multigenerational effects. Notably, genome-wide DNA methylation analyses in mouse embryonic stem cells revealed that genistein perturbed their methylation pattern, showing both regions that were more methylated and regions that were less methylated in genistein-treated cells than in control cells (Sato et al, 2011). In a mouse breast cancer model, maternal genistein supplementation induced both hypoLJ and hyperLJmethylated regions in the genomes of offspring, again pointing to a complex, locusLJspecific action on DNA methylation (Chen et al., 2022).

These findings, together with broader evidence that DNA methylation and histone modifications can mediate multigenerational inheritance of environmental effects, motivated our focus on whether genistein-induced changes in the embryonic environment could have consequences that extend beyond the directly exposed generation.

## Conclusions

While the phenotypic effects of genistein exposure varied in nature and intensity across generations and traits, our findings demonstrate that an environmental exposure during development can induce persistent phenotypic changes, particularly on BW.

Further investigation is needed to understand fully the molecular mechanisms underlying the increased proportion of phenotypic variance explained by the epiline across generations. Although the contribution of genetic drift cannot be fully excluded despite the controlled mating design and the limitation of progeny artificial selection, a molecular analysis would help dissect the potential involvement of non-genetic inheritance mechanisms.

To better disentangle these effects, future studies will focus on dissecting genome-wide epigenetic profiling (e.g., DNA methylation) across generations. For the traits showing epiline effect, EWAS (epigenome wide association studies) and metQTL (methylation quantitative trait loci) analyses should be performed to identify genomic regions where epigenetic marks govern phenotypic changes or where genetic variation is driving epigenetic changes. This will help decipher the variance due to genetic and epigenetic mechanisms and molecular pathways involved in its inheritance.

## Declarations

### Ethics approval and consent to participate

Animals were bred at the UE1295 PEAT (Nouzilly, France, doi: 10.15454/1.5572326250887292E12) with official authorization for the animals (APAFIS#29977-2021010717073072 v5), and PEAT agreement (D371751).

### Consent for publication

Not applicable

### Availability of data and materials

All raw data are available on Zenodo, 10.5281/zenodo.15796146 (https://zenodo.org/records/15796146). The code used to run the analysis is available on GitLab: https://forge.inrae.fr/stacy.rousse/transgenerational_pheno_quails

### Declaration of generative AI and AI-assisted technologies in the writing process

The authors did not resort to using AI or AI-assisted technologies for this work.

## Authors’ contributions

SR performed all the phenotypic data curation, analysis and investigation process, as well as part of the data collection. FP and SL performed all data collection, and participated in the elaboration of the quail design. DG was in charge of the animal facility and contributed to all data collection. LC conceptualized and collected the behavioural data in the open field test. AR contributed to the validation of the statistical method. SL, TZ and FP participated in the project administration, provided resources and secured funding. SE and FP supervised the methodology and the investigation process.

## Acknowledgements

The authors would like to thank Mary Ann Ottinger and Catherine Bennetau-Pelissero for their advice and help concerning genistein. Many thanks to all the collaborators and partners, in particular Ingrid David for her statistical insights and the colleagues from PEAT, especially David Gourichon, Sandrine Rivière and Michael Troquet who carefully reared the quails and kindly helped for all experiments. Finally, special thanks to the Genobioinfo platform for data management. This work has been submitted as a pre-print: https://doi.org/10.1101/2025.07.04.663137

## Additional files

### Additional file 1 Table S1

Format: xls

Title: Summary statistics for all phenotypic traits in different generations

Description: Nobs = number of phenotypic records; mean; sd = standard deviation; CV = coefficient of variation; min = minimum; max = maximum; ^1^BW = body weights; ^2^TW = tissue weights. All tissue weights are adjusted (corrected by the weight at slaughter); ^3^PROD = production traits. Egg weight is adjusted (corrected by the weight at slaughter); ^4^BHV = behavioural traits. Results to an isolation test; ^5^abdominal fat; ^6^frequency at periphery.

### Additional file 2 Figure S1

Format: pdf

Title: The evolution of Body Weights per epiline and sex

Description: The longitudinal evolution of the average BW with the age of the quail (expressed in weeks) is displayed per sex and epiline, in each generation. The solid line links the average BW of epi-quails and the dashed line the epi+. Differences between epilines become visible in Generation 1 in females and in Generation 2 in males when animals reach adult age (7 weeks).

### Additional file 3 Figure S2

Format: tiff

Title: Differences between epilines in adjusted tissue weights

Description: All tissue weights are adjusted by the adult body weight. The study was conducted for each combination of sex and generation independently. N values indicate sample sizes within each sex and epiline per generation. The asterisk indicates the level of significance of the Kruskal-Wallis test results, ns p-value > 0.05 (non significant), * p-value < 0.05, ** p-value < 0.01, *** p-value < 0.001 and **** p-value < 1e^-4^.

Overall, no significant effect of the epiline is detected, except for adjusted heart weight in G1 for both sexes. The significant epiline effect detected in G2 males for this trait are most likely due to an outlier individual in epi-. Trait average was consistent between generations, apart from a slight decrease in adjusted liver weight from G0 to G2.

### Additional file 4 Figure S3

Format: tiff

Title: Differences between epilines in production trait

Description: Egg weight is adjusted by the adult body weight. The study was conducted for each generation independently. N values indicate sample sizes within each sex and epiline per generation. The asterisk indicates the level of significance of the Kruskal-Wallis test results, ns p-value > 0.05 (non significant), * p-value < 0.05, ** p-value < 0.01, *** p-value < 0.001 and **** p-value < 1e^-4^.

Overall, no significant effect of the epiline was detected, except for adjusted egg weight in G0. The significant effect detected for precocity in G1 is most probably due to individual singularities. Trait average was consistent between generations.

### Additional file 5 Figure S4

Format: tiff

Title: Differences between epilines in behavioural traits

Description: Total distance is expressed in millimetres. Duration at centre corresponds to the time spent in the inner area, expressed in seconds. The frequency at the periphery corresponds to the number of times the animal stepped out of the inner area. N values indicate sample sizes within each sex and epiline per generation. The asterisk indicates the level of significance of the Kruskal-Wallis test results, ns p-value > 0.05 (non significant), * p-value < 0.05, ** p-value < 0.01, *** p-value < 0.001 and **** p-value < 1e^-4^.

Overall, no significant effect of the epiline was detected, except for the duration spent at the centre of the arena in G1 males. The significant effect detected in G2 males for the frequency at the periphery is most probably due to an individual outlier in the control epiline (epi-).

### Additional file 6

Format: pdf

Title: Phenotypic raw and transformed distributions.

**A.** Summary table of applied transformations for normalization and skewness values. The different values of skewness of the data are reported in this table. The objective is to apply the transformation that minimizes the skewness to meet the assumptions of normality of the linear model applied. Raw: raw data, no transformation applied. log10, sqrt, inv refer to Log10, Square root and inverse transformations respectively. The ‘best’ column summarises the choice of transformation to apply for the trait considered. For body weight traits (BW), BW at one and four weeks were not transformed, while a square root transformation was applied for BW at seven weeks and at slaughter. **B.** Phenotypic distributions in G0; **C.** Phenotypic distributions in G1; **D.** Phenotypic distributions in G2. All phenotypic traits collected in each generation are represented in the graphs (black distributions). When a transformation was chosen to minimize skewness (see table A.), the distribution of the transformed trait is represented (purple distributions, ‘_trans’ suffix). **E-J.** Residuals Q-Q plots from raw phenotypes before applying Kruskal-Wallis tests respectively in G0 males, G0 females, G1 males, G1 females, G2 males and G2 females. The results are shown per epiline to compare trends within each group. **K**. Summary table of Shapiro-Wilk normality tests in each sex and generation. The Null Hypothesis tested is H0 “The distribution of the data does not deviate from a normal distribution” at a significance threshold of 0.05. **L**. Summary table of Kruskal-Wallis mean comparisons between epilines for each sex and generation. The Null Hypothesis tested is H0 “There is no difference between epilines” at a significance threshold of 0.05.

### Additional file 7

Format: pdf

Title: Contribution of parameters to phenotypic variability per generation.

**A.** Summary table of the linear model fitted for each trait with the best set of parameters kept after the stepAIC. The ‘_trans’ suffix was added to each trait when a transformation was applied to minimize skewness (see Additional file 6). The interaction parameters considered are detailed in the linear model equation. **B.** Contribution of parameters (sex, family, epiline, reproductive status and their interactions) to phenotypic variability (in percentage) per generation. Portion of phenotypic variability explained by the effect of sex, family, epiline, reproductive status and their interaction as extracted from the best linear model for adjusted tissue weights, production traits and behaviours. The reproductive status (repro status, yellow) clarifies whether the quail was used as a parent for the next generation. This status is not considered for the analysis of BW because the mating plan is elaborated after weighing at one, four and seven weeks and therefore this variable was not relevant for the trait. The interaction parameter was divided into two categories: the ‘interaction epi’ parameter gathers sex:epiline, epiline:family and epiline:repro while ‘interaction non epi’ gathers sex:family, sex:repro and family:repro. Overall, the epiline (pink) shows no effect, or negligible given the substantial amount of residual variance. The incapacity of the model in capturing a solid portion of the trait variability questions the confidence in the estimates of the effects obtained. **C.** Summary table of the estimates of the contribution of parameters (and standard deviation) to phenotypic variability (expressed in percentage). The effect of epiline, family, sex, reproductive status, interaction epi and interaction non epi and residuals of each trait are expressed in percentage with the standard deviation in brackets. The estimates were calculated from the extraction of the sum of squares from analyses of variance (ANOVA type III) results. All figures are percentages and represent the average contribution of the different factors to phenotypic variability using the estimates obtained across the 100 folds of cross validations. The interaction parameter was divided into two categories: the ‘interaction epi’ parameter gathers sex:epiline, epiline:family and epiline:repro while ‘interaction non epi’ gathers sex:family, sex:repro and family:repro. **D.** Model robustness (R²) and correlations after cross validations. The coefficient of determination R² (blue) and the correlation between observed and predicted values in the Test set (yellow) were determined in 100 cross validations, to evaluate the predictive performance of linear models. The results show the distribution of R² and correlation values in each of the 100 cross validations, and are displayed per generation (x-axis) for each phenotype.

### Additional file 8

Format: pdf

Title: Simulation of a phenotype resembling body weight at slaughter over 5 generations.

The phenotypic distributions in two epilines are simulated in each generation after a similar mating design as the one explored in our study. The trait was simulated as a polygenic trait (1000 QTL for 10000 chromosome segments) of mean 200, to resemble body weight at slaughter, variance 50 and an environmental variance of heritability 0.5 (as described in Karami et al. 2025). An initial founder population (PM) was simulated, it comprised 40 individuals, 20 males and 20 females, crossed with each other and each producing 10 progenies. These crosses initiated 20 founder families, as in our experimental design. From their offspring, 20 males and 40 females were selected, each male was mated with two females, therefore initiating the epilines (EPIP and EPIM). The males and females to mate were chosen to be from different founder families. Thereafter, mating was performed within each epiline independently and in mirror with the aim to cross different families in every generation. The simulation framework was repeated 100 times independently. The simulations were made using AlphaSimR (Faux et al. 2016, Gaynor et al. 2021)

**A.** Violin plot showing the distribution of the differences (in absolute) in phenotypic values between the two epilines, for each sex. The black dots are the 100 simulated values, the red dots are the values obtained from real data.

Results display differences between epilines in phenotypic values for a number of simulations and for all generations. This supports the assumption made by David & Ricard 2023, that the difference in phenotypes can happen due to drift only. However, the differences in body weight at slaughter between epilines obtained with our simulations are lower than the ones observed with the real data, from G1 onwards in females, and in G2 in males, which concurs with our conclusions on the potential epiline effect on this trait.

