## Supplementary table 1 for "Transmission of an environmental modification in quails across three generations: changes in the contribution to phenotypic variability"

**Supplementary Table S1** Descriptive statistics of phenotypic records for individuals of the G-1 generation.

| Trait | Nobs | mean | sd | CV | min | max |
|---|---|---|---|---|---|---|
| BW[1] 1 week | 223 | 30.78 | 3.22 | 0.1 | 12 | 39 |
| BW 4 weeks | 223 | 126.59 | 10.1 | 0.08 | 78 | 150 |
| BW 7 weeks | 223 | 153.92 | 10.72 | 0.07 | 122 | 181 |
| BW slaughter | 128 | 218.66 | 26.63 | 0.12 | 164 | 269 |
| abdo.[2] fat weight | 117 | 0.01 | 0.01 | 0.52 | 0 | 0.02 |
| total distance | 209 | 1243.2 | 1386.63 | 1.12 | 59.97 | 8038.63 |
| duration at centre | 205 | 205.6 | 94.85 | 0.46 | 1.84 | 300.08 |
| freq.[3] at periphery | 209 | 10.26 | 13.29 | 1.29 | 0 | 69 |

Nobs = number of phenotypic records; mean; sd = standard deviation; CV = coefficient of variation; min = minimum; max = maximum.

[1] BW: body weight

[2] Abdominal fat weight.

[3] Frequency at periphery
