## Supplementary figures and images for "Transmission of an environmental modification in quails across three generations: changes in the contribution to phenotypic variability"

### Supplementary figure 1

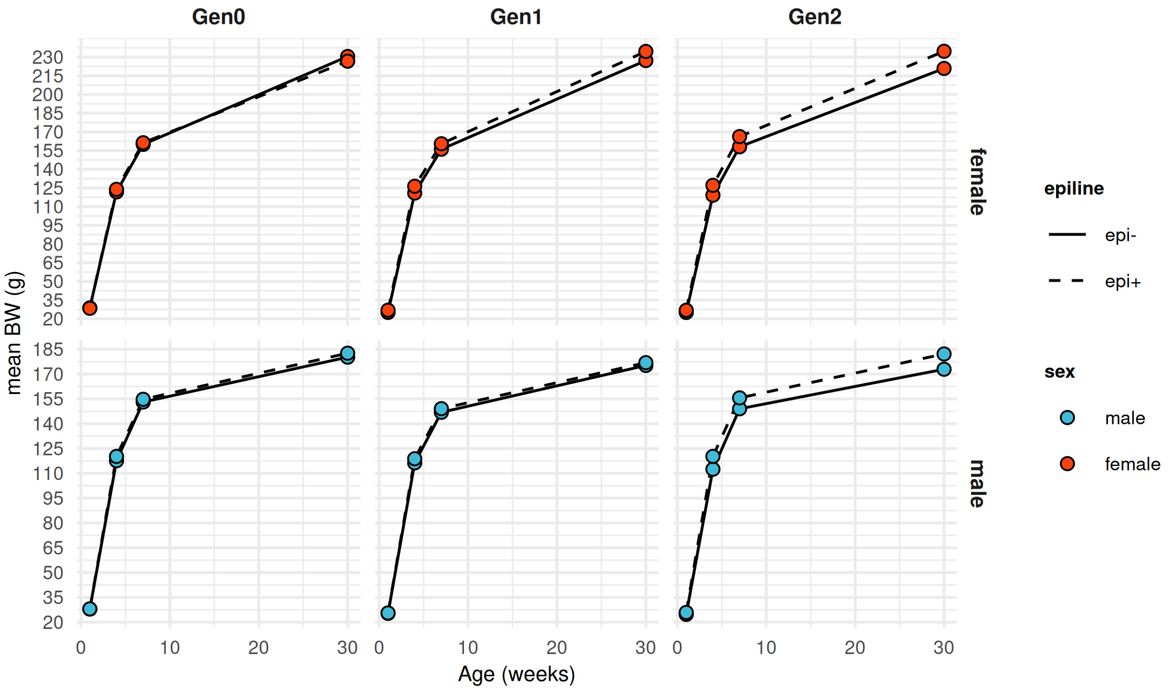

### Supplementary figure 2

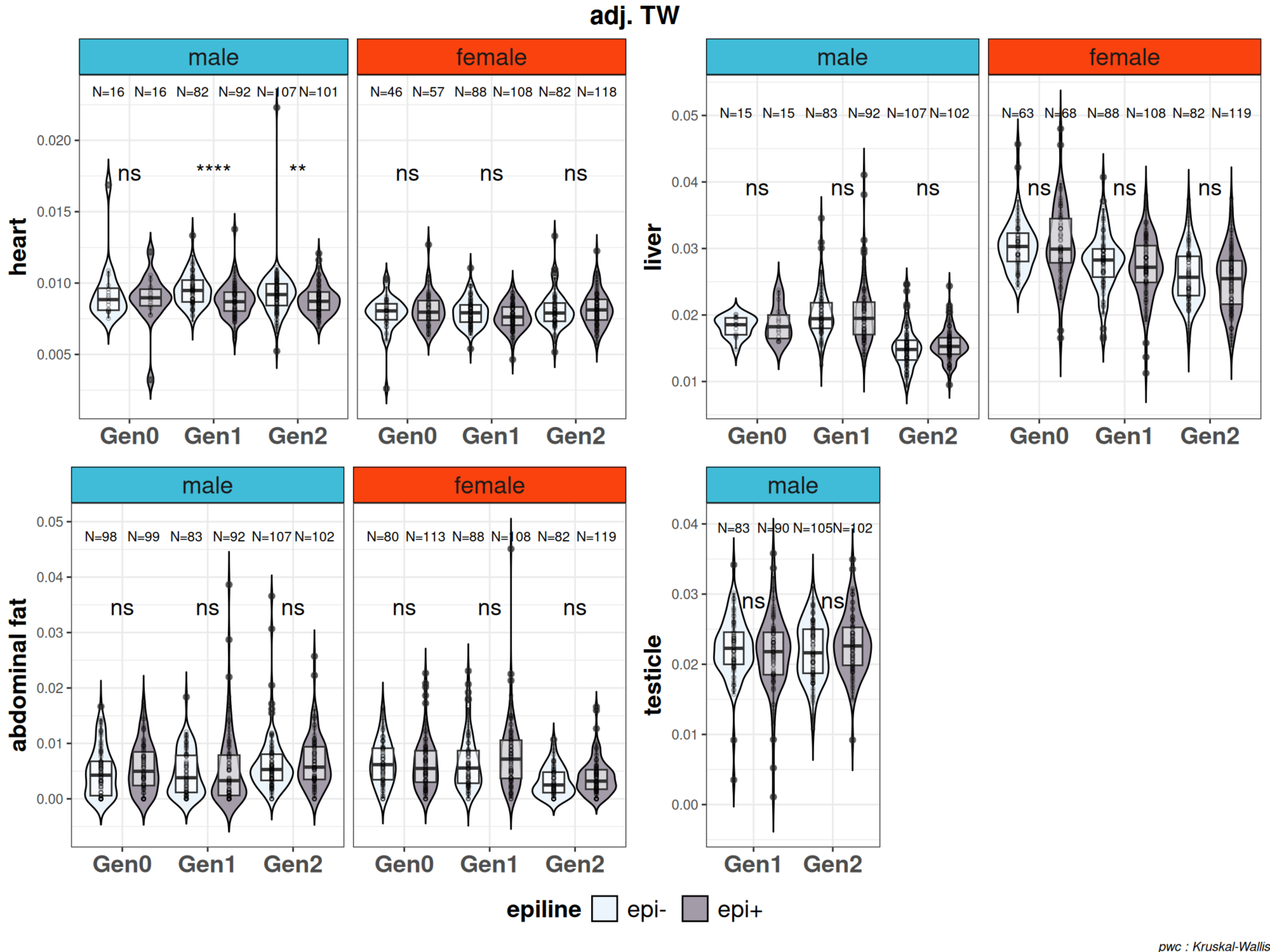

### Supplementary figure 3

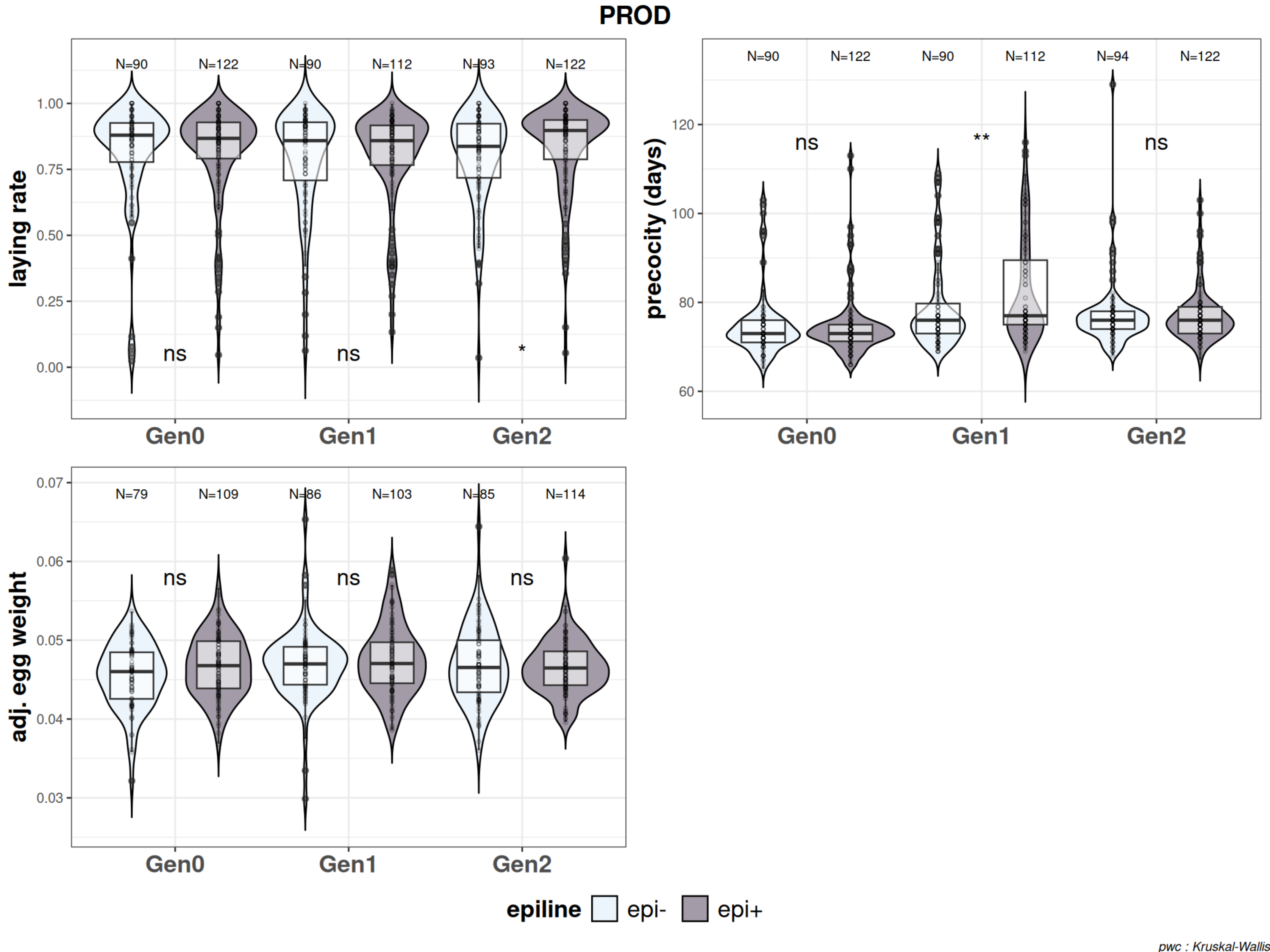

### Supplementary figure 4

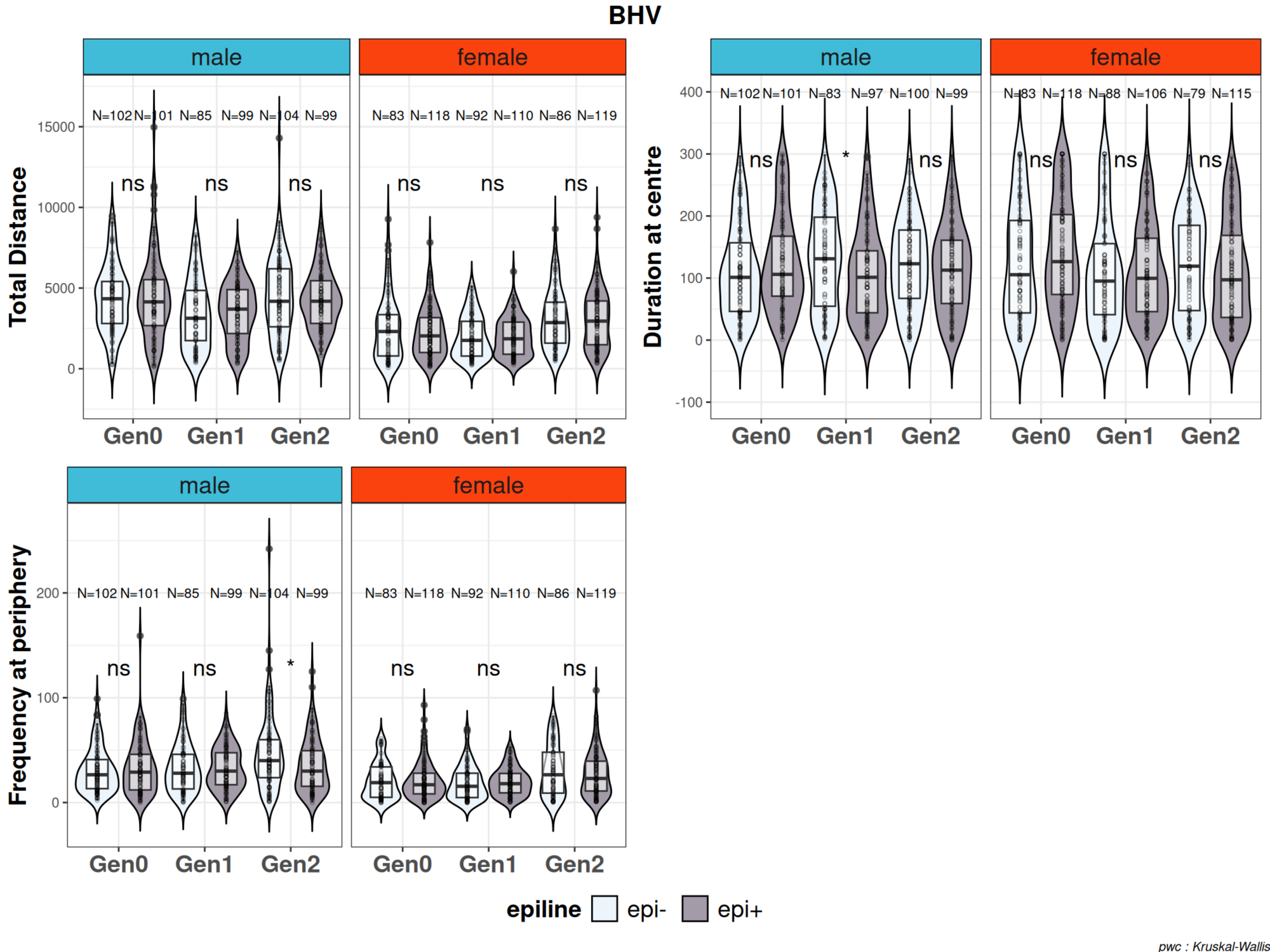
