## Supplementary material 6 for "Transmission of an environmental modification in quails across three generations: changes in the contribution to phenotypic variability"

**Additional file 6:** Distribution of raw and transformed phenotypes. Distributions in black highlight the raw trait distributions (without normalization). The distributions in purple are normalized trait distributions after transformation. The "trans" suffix was added to highlight that the trait was transformed. All phenotypic traits collected at each generation are represented. bw_1w: bodyweight at one week, bw_4w: bodyweight at four weeks, bw_7w: bodyweight at seven weeks, bw_sl: bodyweight at slaughter, adj_heartW: adjusted heart weight, adj_livrW: adjusted liver weight, adj_adipW: adjusted abdominal fat weight, LR: laying rate, adj_EW: adjusted egg weight, totalDist: total distance, centreDuration: duration at centre, periphFreq: frequency at periphery.
**A.** Summary table of applied transformations for normalization and skewness values. raw: no transformation, log10: Log10 transformation, sqrt: square root transformation, inv: inverse transformation, best: transformation chosen that minimizes data skewness. **B.** Phenotypic distributions in G0. **C.** Phenotypic distributions in G1. **D.** Phenotypic distributions in G2. **E-J.** Residuals Q-Q plots from raw phenotypes before ANOVA respectively in G0 males, females, G1 males, females, G2 males, females. **K.** Summary table of Shapiro-Wilk normality tests in each sex and generation. The Null Hypothesis tested is H0 "The distribution of the data does not deviate from a normal distribution" at a significance threshold of 0.05. **L.** Summary table of Kruskal-Wallis mean comparisons between epilines for each sex and generation. The Null Hypothesis tested is H0 "There is no difference between epilines" at a significance threshold of 0.05.

# A.

| **Trait** | **raw** | **log10** | **sqrt** | **inv** | **best** |
|---|---|---|---|---|---|
| bw_1w | 0.010 | 0.483 | 0.219 | 1.211 | raw |
| bw_4w | 0.071 | 0.373 | 0.218 | 0.717 | raw |
| bw_7w | 0.151 | 0.051 | 0.050 | 0.256 | sqrt |
| bw_sl | 0.109 | 0.112 | 0.0002 | 0.347 | sqrt |
| adj_heartW | 1.552 | 0.382 | 0.522 | 3.241 | log10 |
| adj_livrW | 0.250 | 0.322 | 0.038 | 0.953 | sqrt |
| adj_adipW | 1.572 | 1.535 | 0.159 | 1.500 | sqrt |
| adj_testisW | 0.438 | 4.472 | 1.607 | 15.976 | raw |
| LR | 1.916 | 4.408 | 2.767 | 10.605 | raw |
| precocity | 2.137 | 1.794 | 1.961 | 1.481 | inv |
| adj_EW | 0.245 | 0.248 | 0.003 | 0.802 | sqrt |
| totalDist | 0.910 | 1.071 | 0.012 | 4.437 | sqrt |
| centreDuration | 0.374 | 2.085 | 0.323 | 21.861 | sqrt |

| | | | | | |
|---|---|---|---|---|---|
| periphFreq | 1.691 | 0.853 | 0.107 | 2.886 | sqrt |

**B.**

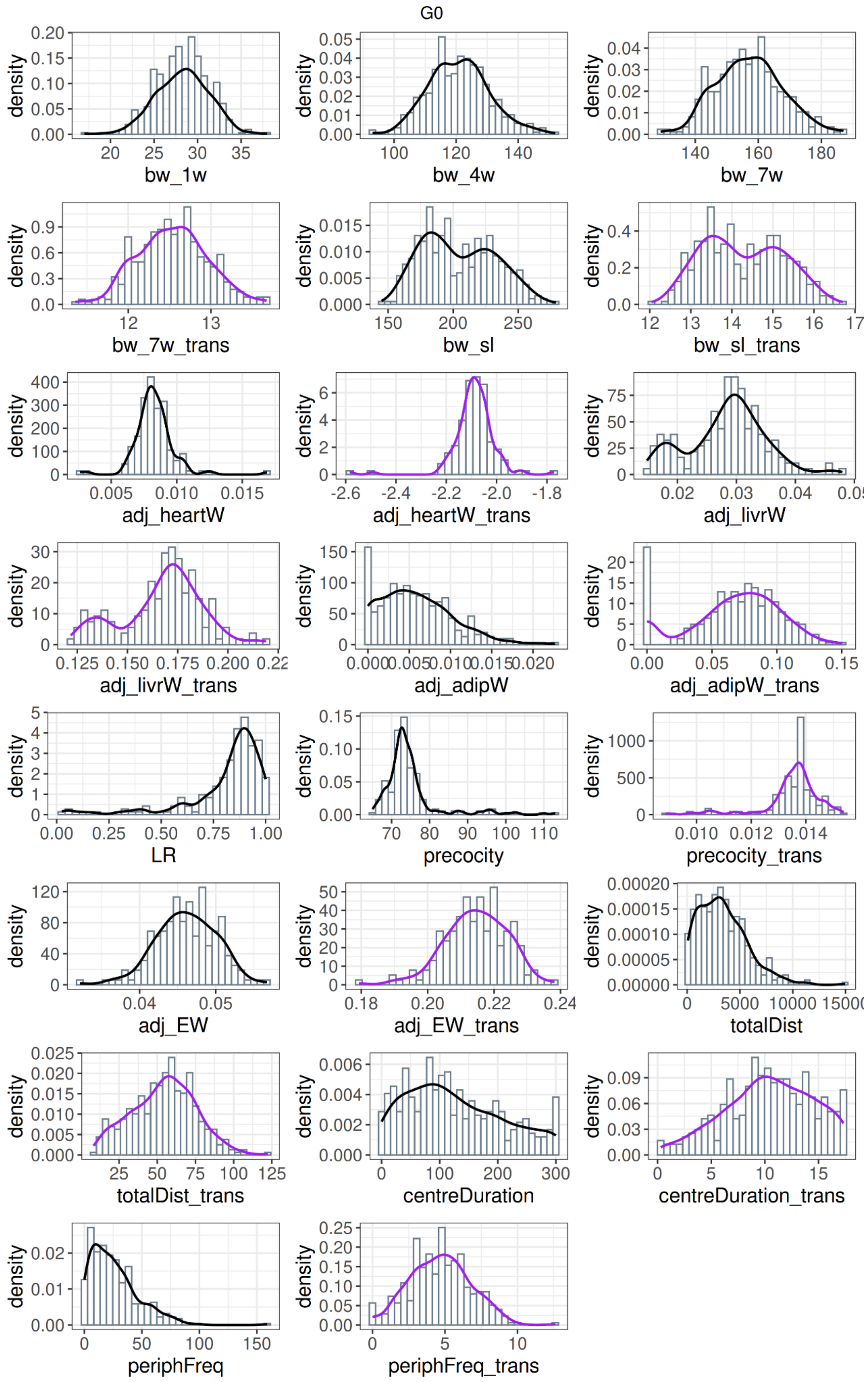

C.

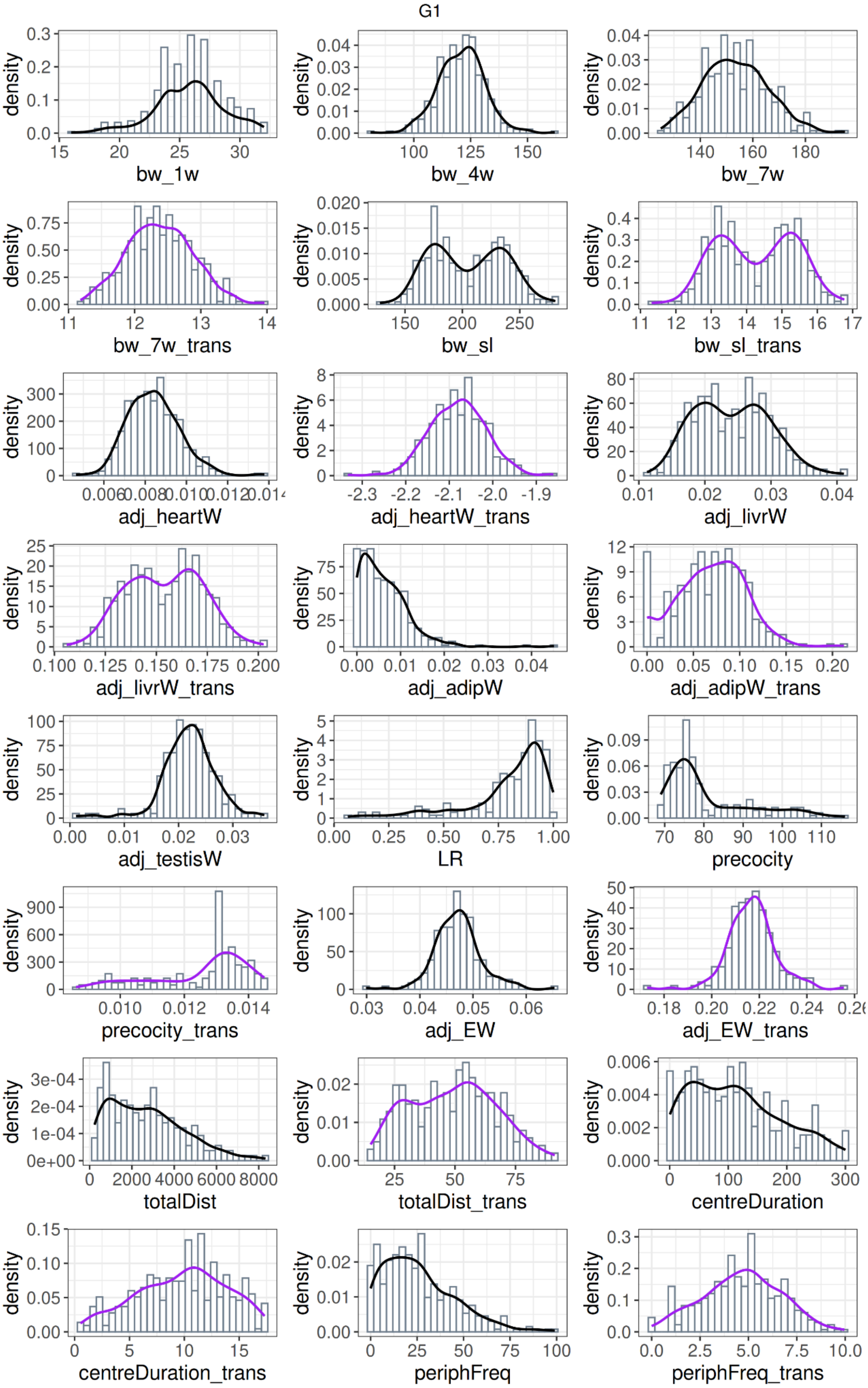
G1
density
bw_1w
bw_4w
bw_7w
bw_7w_trans
bw_sl
bw_sl_trans
adj_heartW
adj_heartW_trans
adj_livrW
adj_livrW_trans
adj_adipW
adj_adipW_trans
adj_testisW
LR
precocity
precocity_trans
adj_EW
adj_EW_trans
totalDist
totalDist_trans
centreDuration
centreDuration_trans
periphFreq
periphFreq_trans

**D.**

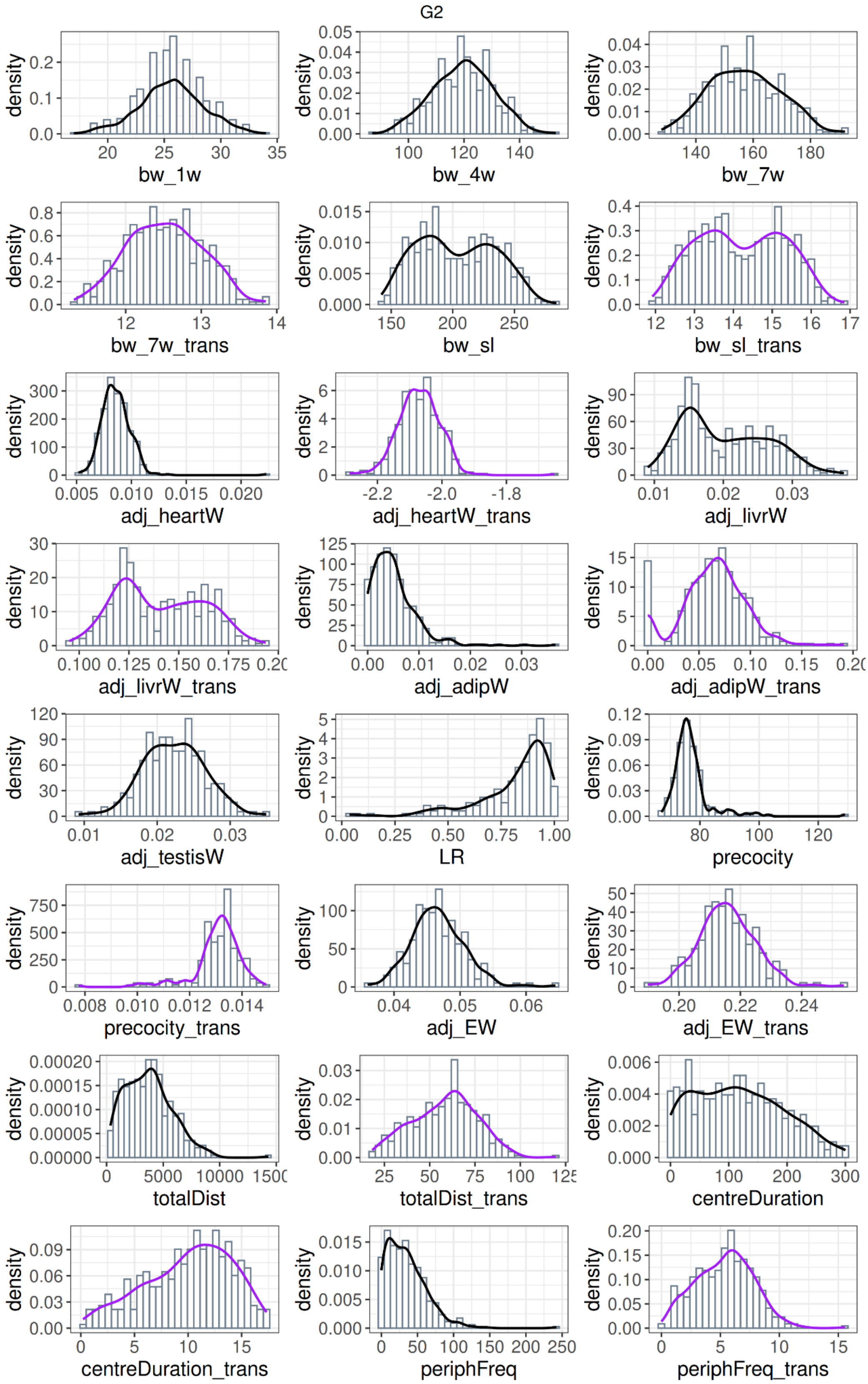

G2
density
bw_1w
bw_4w
bw_7w
bw_7w_trans
bw_sl
bw_sl_trans
adj_heartW
adj_heartW_trans
adj_livrW
adj_livrW_trans
adj_adipW
adj_adipW_trans
adj_testisW
LR
precocity
precocity_trans
adj_EW
adj_EW_trans
totalDist
totalDist_trans
centreDuration
centreDuration_trans
periphFreq
periphFreq_trans

**E.**

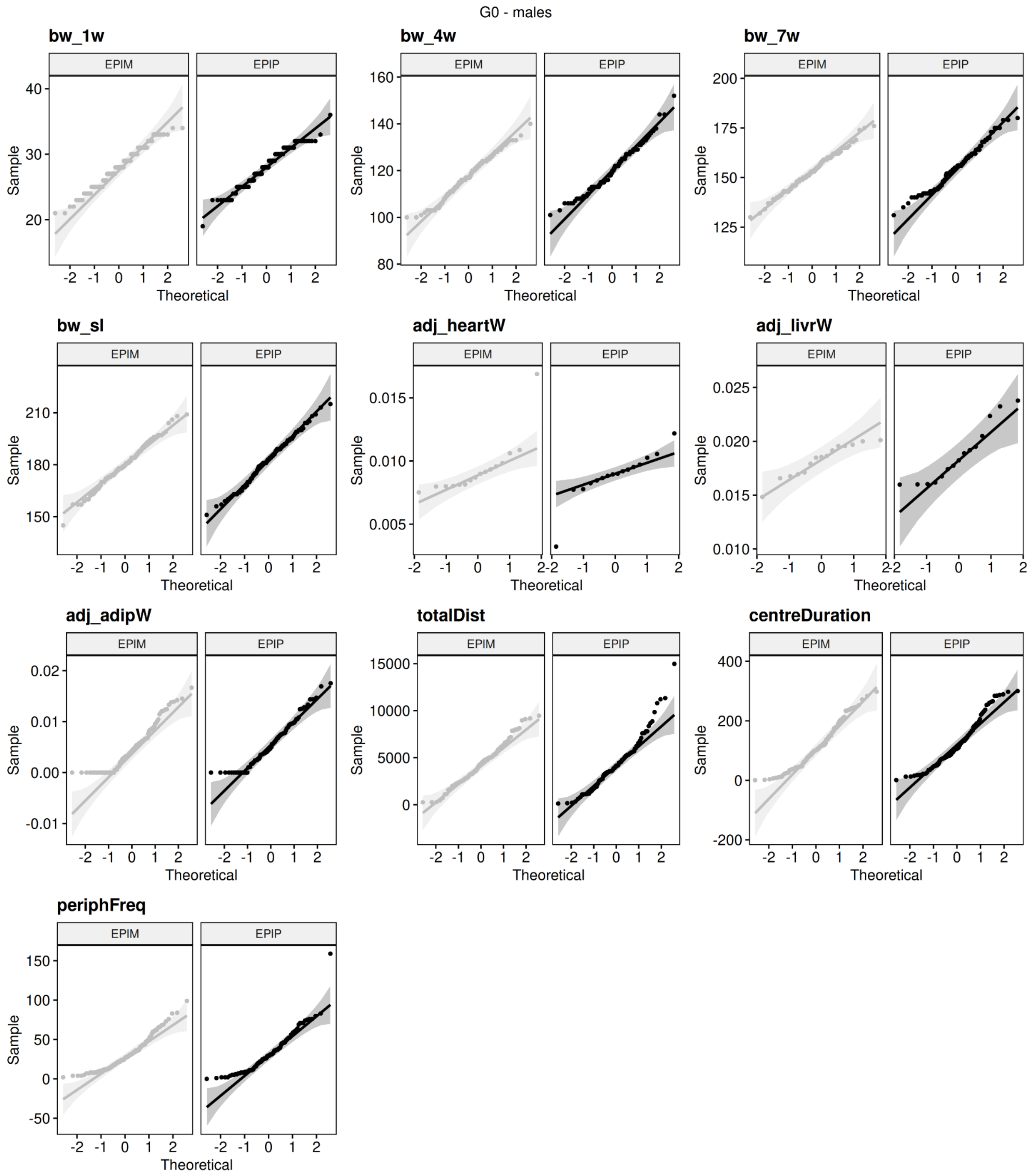

G0 - males
bw_1w
bw_4w
bw_7w
bw_sl
adj_heartW
adj_livrW
adj_adipW
totalDist
centreDuration
periphFreq
EPIM
EPIP
Sample
Theoretical

F.

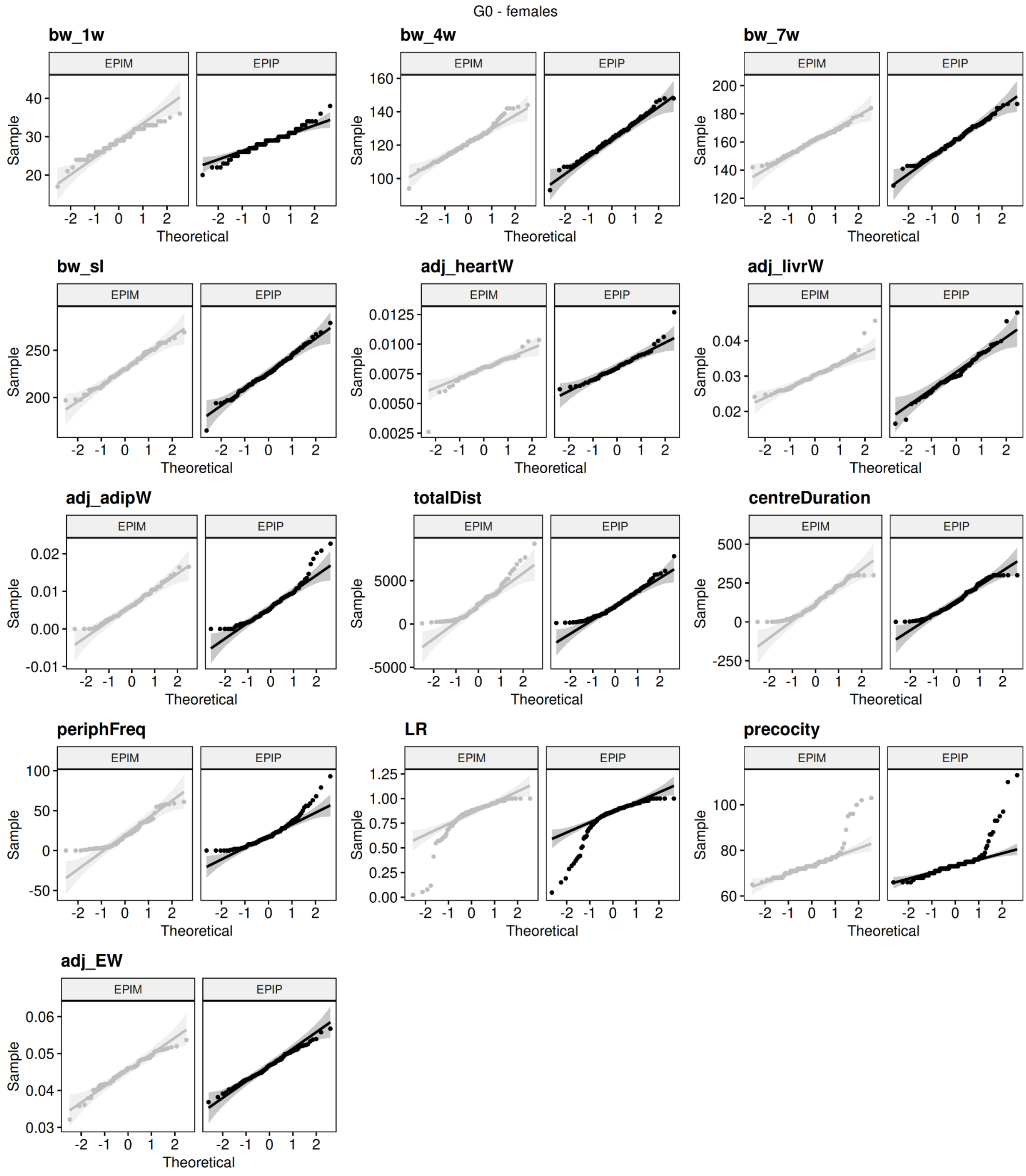

G0 - females
bw_1w
bw_4w
bw_7w
bw_sl
adj_heartW
adj_livrW
adj_adipW
totalDist
centreDuration
periphFreq
LR
precocity
adj_EW
EPIM
EPIP
Sample
Theoretical

G.

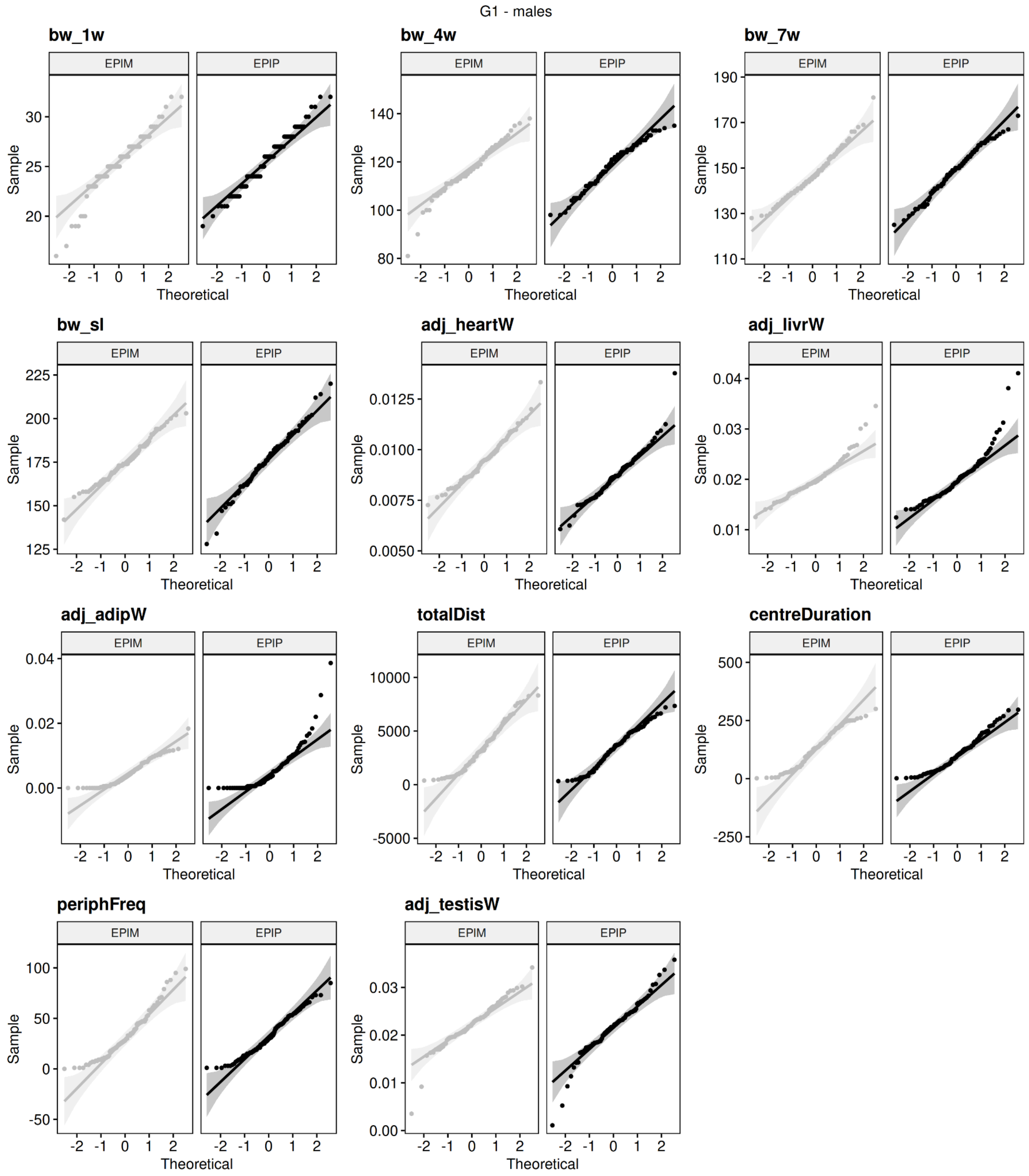

G1 - males
bw_1w
bw_4w
bw_7w
bw_sl
adj_heartW
adj_livrW
adj_adipW
totalDist
centreDuration
periphFreq
adj_testisW
EPIM
EPIP
Sample
Theoretical

H.

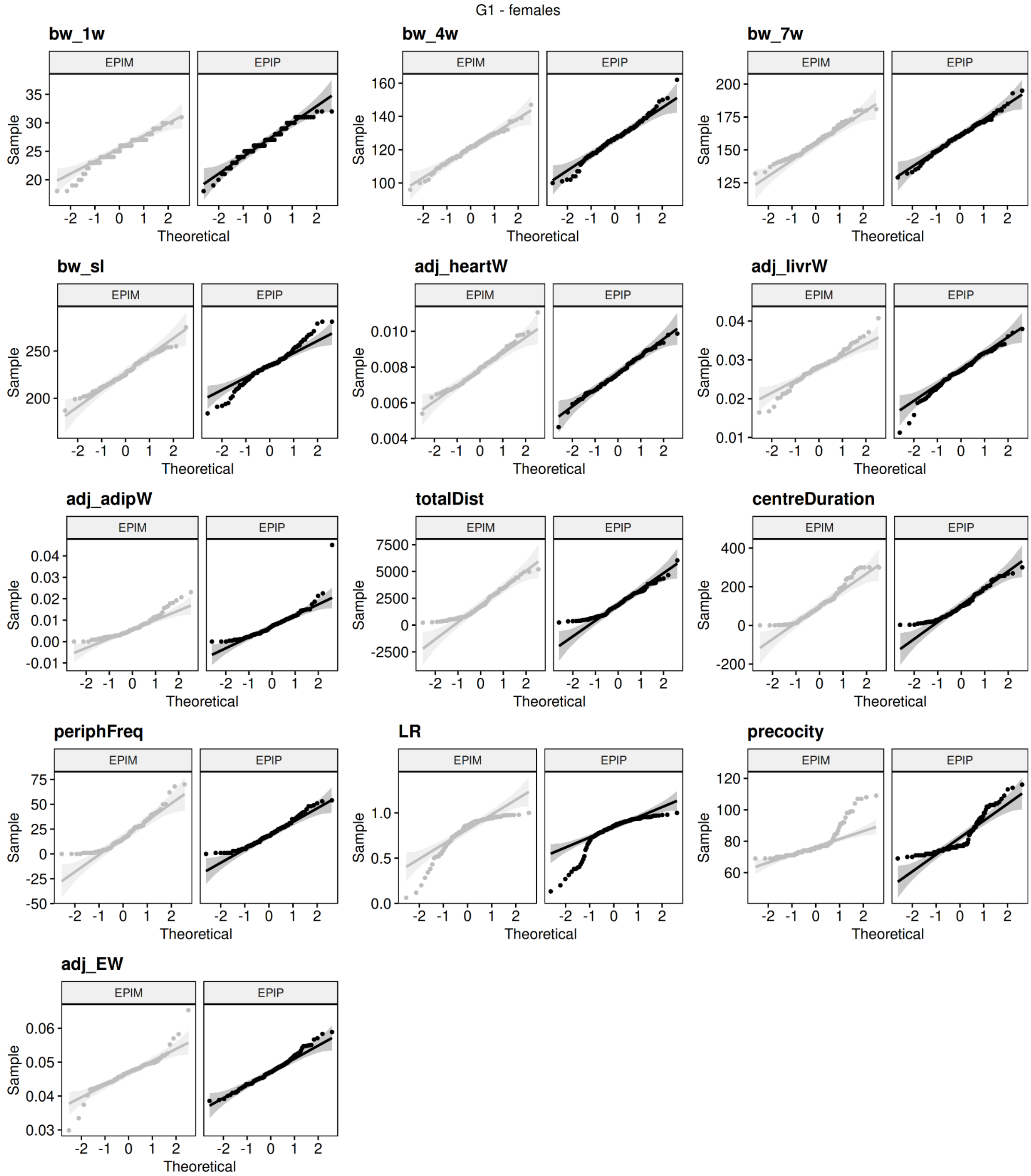
G1 - females
bw_1w
bw_4w
bw_7w
bw_sl
adj_heartW
adj_livrW
adj_adipW
totalDist
centreDuration
periphFreq
LR
precocity
adj_EW
EPIM
EPIP
Sample
Theoretical

I.

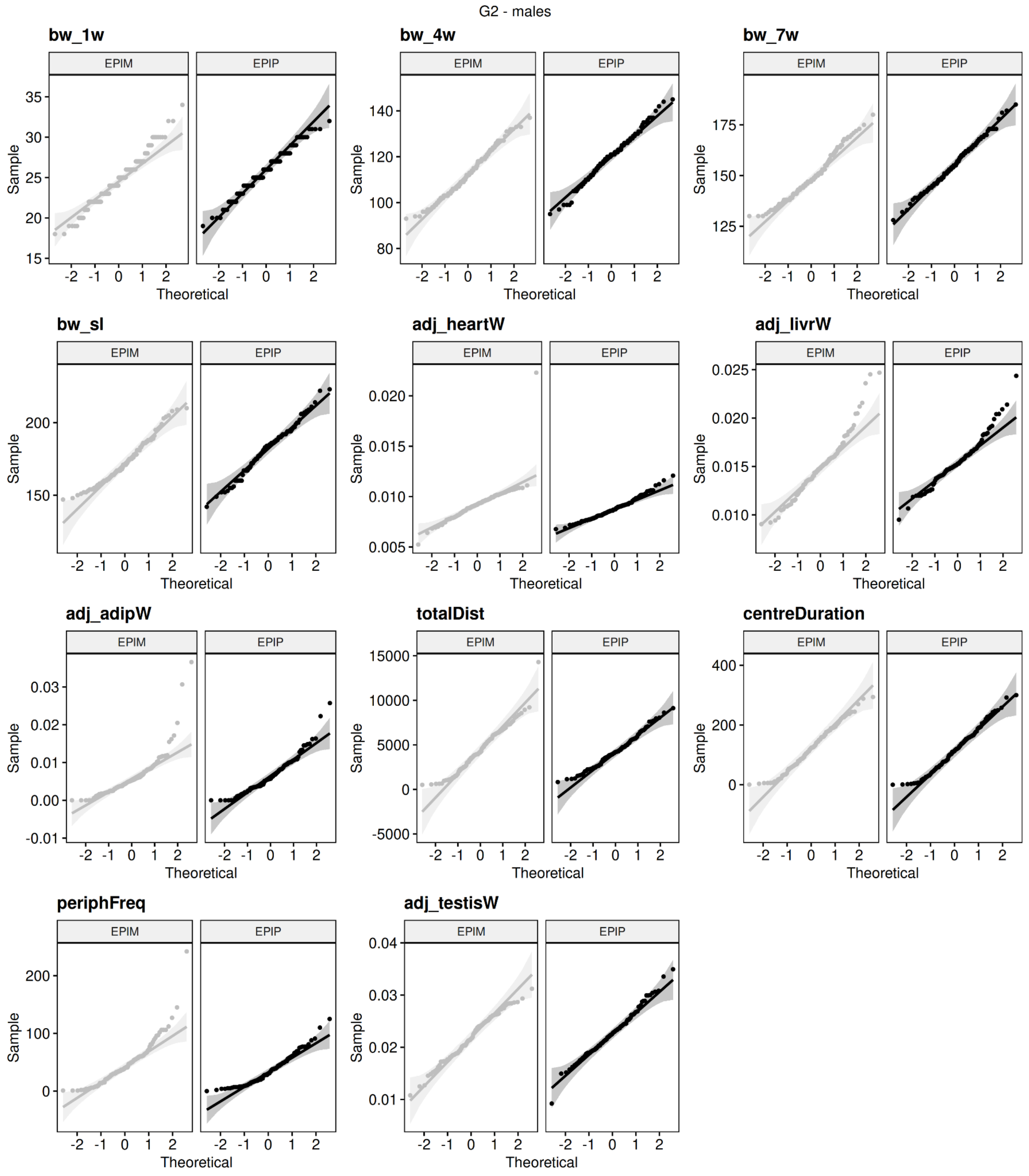
G2 - males
bw_1w
bw_4w
bw_7w
bw_sl
adj_heartW
adj_livrW
adj_adipW
totalDist
centreDuration
periphFreq
adj_testisW
EPIM
EPIP
Sample
Theoretical

**J.**

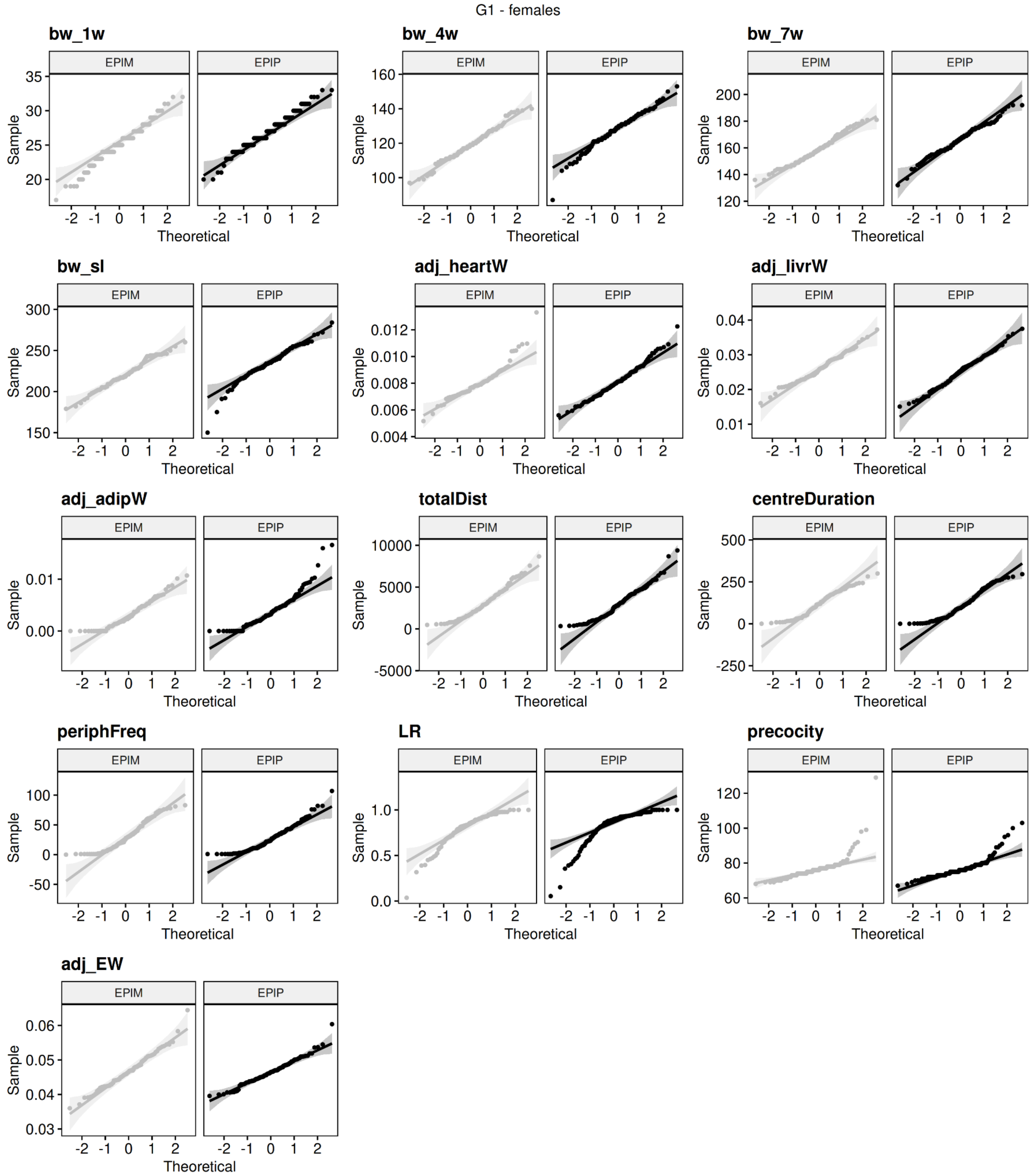
G1 - females
bw_1w
bw_4w
bw_7w
bw_sl
adj_heartW
adj_livrW
adj_adipW
totalDist
centreDuration
periphFreq
LR
precocity
adj_EW
EPIM
EPIP
Sample
Theoretical

**K.**

| Trait | generation | sex | pval | test |
|---|---|---|---|---|
| bw_1w | Gen0 | male | 0.0055 | 0.9814 |
| bw_4w | Gen0 | male | 0.05556 | 0.98766 |
| bw_7w | Gen0 | male | 0.14311 | 0.99018 |
| bw_sl | Gen0 | male | 0.92455 | 0.99644 |
| adj_heartW | Gen0 | male | 0.00004 | 0.79872 |
| adj_livrW | Gen0 | male | 0.16773 | 0.94987 |
| adj_adipW | Gen0 | male | 0 | 0.93188 |
| centreDuration | Gen0 | male | 0 | 0.95049 |
| periphFreq | Gen0 | male | 0 | 0.90406 |
| totalDist | Gen0 | male | 0.00003 | 0.96234 |
| bw_1w | Gen0 | female | 0.01912 | 0.98443 |
| bw_4w | Gen0 | female | 0.14607 | 0.98999 |
| bw_7w | Gen0 | female | 0.35133 | 0.99248 |
| bw_sl | Gen0 | female | 0.51931 | 0.99332 |
| adj_heartW | Gen0 | female | 0.00004 | 0.93073 |
| adj_livrW | Gen0 | female | 0.00131 | 0.96326 |
| adj_adipW | Gen0 | female | 0 | 0.94621 |
| centreDuration | Gen0 | female | 0 | 0.94859 |
| periphFreq | Gen0 | female | 0 | 0.90435 |
| totalDist | Gen0 | female | 0 | 0.92232 |
| LR | Gen0 | female | 0 | 0.74957 |
| adj_EW | Gen0 | female | 0.57197 | 0.99346 |
| precocity | Gen0 | female | 0 | 0.70465 |
| bw_1w | Gen1 | male | 0.00356 | 0.9769 |

| | | | | |
|---|---|---|---|---|
| bw_4w | Gen1 | male | 0.01128 | 0.98078 |
| bw_7w | Gen1 | male | 0.5876 | 0.99352 |
| bw_sl | Gen1 | male | 0.34441 | 0.99112 |
| adj_heartW | Gen1 | male | 0.00354 | 0.97542 |
| adj_livrW | Gen1 | male | 0 | 0.89834 |
| adj_adipW | Gen1 | male | 0 | 0.80282 |
| adj_testisW | Gen1 | male | 0.00001 | 0.94869 |
| centreDuration | Gen1 | male | 0.00003 | 0.95721 |
| periphFreq | Gen1 | male | 0.00004 | 0.96003 |
| totalDist | Gen1 | male | 0.00091 | 0.97188 |
| bw_1w | Gen1 | female | 0.0003 | 0.97109 |
| bw_4w | Gen1 | female | 0.10728 | 0.98881 |
| bw_7w | Gen1 | female | 0.60532 | 0.99418 |
| bw_sl | Gen1 | female | 0.58107 | 0.99376 |
| adj_heartW | Gen1 | female | 0.49325 | 0.99313 |
| adj_livrW | Gen1 | female | 0.03181 | 0.98471 |
| adj_adipW | Gen1 | female | 0 | 0.86057 |
| centreDuration | Gen1 | female | 0 | 0.93779 |
| periphFreq | Gen1 | female | 0 | 0.93923 |
| totalDist | Gen1 | female | 0 | 0.9475 |
| LR | Gen1 | female | 0 | 0.8038 |
| adj_EW | Gen1 | female | 0.00018 | 0.96668 |
| precocity | Gen1 | female | 0 | 0.81536 |
| bw_1w | Gen2 | male | 0.0136 | 0.98674 |
| bw_4w | Gen2 | male | 0.24305 | 0.99308 |
| bw_7w | Gen2 | male | 0.01044 | 0.98605 |
| bw_sl | Gen2 | male | 0.05579 | 0.98714 |

| | | | | |
|---|---|---|---|---|
| adj_heartW | Gen2 | male | 0 | 0.76497 |
| adj_livrW | Gen2 | male | 0.00004 | 0.96431 |
| adj_adipW | Gen2 | male | 0 | 0.83689 |
| adj_testisW | Gen2 | male | 0.95259 | 0.99685 |
| centreDuration | Gen2 | male | 0.00274 | 0.97748 |
| periphFreq | Gen2 | male | 0 | 0.87496 |
| totalDist | Gen2 | male | 0.0004 | 0.97161 |
| bw_1w | Gen2 | female | 0.00247 | 0.98148 |
| bw_4w | Gen2 | female | 0.53771 | 0.99457 |
| bw_7w | Gen2 | female | 0.23206 | 0.99217 |
| bw_sl | Gen2 | female | 0.08 | 0.98793 |
| adj_heartW | Gen2 | female | 0.00006 | 0.96436 |
| adj_livrW | Gen2 | female | 0.69729 | 0.99467 |
| adj_adipW | Gen2 | female | 0 | 0.90878 |
| centreDuration | Gen2 | female | 0 | 0.94936 |
| periphFreq | Gen2 | female | 0 | 0.92953 |
| totalDist | Gen2 | female | 0 | 0.95454 |
| LR | Gen2 | female | 0 | 0.81864 |
| adj_EW | Gen2 | female | 0.00183 | 0.97618 |
| precocity | Gen2 | female | 0 | 0.72279 |

**L.**

| Trait | generation | sex | pval | test |
|---|---|---|---|---|
| bw_1w | Gen0 | male | 0.68216 | 0.1677 |
| bw_4w | Gen0 | male | 0.09981 | 2.70853 |
| bw_7w | Gen0 | male | 0.33577 | 0.92652 |
| bw_sl | Gen0 | male | 0.17709 | 1.82187 |
| adj_heartW | Gen0 | male | 0.93991 | 0.00568 |
| adj_livrW | Gen0 | male | 0.96691 | 0.00172 |
| adj_adipW | Gen0 | male | 0.16464 | 1.9311 |
| centreDuration | Gen0 | male | 0.25636 | 1.2883 |
| periphFreq | Gen0 | male | 0.58415 | 0.29957 |
| totalDist | Gen0 | male | 0.80374 | 0.06176 |
| bw_1w | Gen0 | female | 0.35732 | 0.84729 |
| bw_4w | Gen0 | female | 0.09673 | 2.75863 |
| bw_7w | Gen0 | female | 0.41576 | 0.66227 |
| bw_sl | Gen0 | female | 0.12459 | 2.35866 |
| adj_heartW | Gen0 | female | 0.64952 | 0.2065 |
| adj_livrW | Gen0 | female | 0.73145 | 0.11778 |
| adj_adipW | Gen0 | female | 0.37307 | 0.79343 |
| centreDuration | Gen0 | female | 0.26985 | 1.21749 |
| periphFreq | Gen0 | female | 0.83123 | 0.04542 |
| totalDist | Gen0 | female | 0.83226 | 0.04486 |
| LR | Gen0 | female | 0.86775 | 0.02773 |
| adj_EW | Gen0 | female | 0.10298 | 2.65884 |
| precocity | Gen0 | female | 0.78207 | 0.07652 |
| bw_1w | Gen1 | male | 0.86248 | 0.03001 |
| bw_4w | Gen1 | male | 0.05526 | 3.67414 |

| | | | | |
|---|---|---|---|---|
| bw_7w | Gen1 | male | 0.05538 | 3.67046 |
| bw_sl | Gen1 | male | 0.30982 | 1.03143 |
| adj_heartW | Gen1 | male | 0.00006 | 16.22683 |
| adj_livrW | Gen1 | male | 0.6167 | 0.25053 |
| adj_adipW | Gen1 | male | 0.80153 | 0.06319 |
| adj_testisW | Gen1 | male | 0.40768 | 0.68557 |
| centreDuration | Gen1 | male | 0.03149 | 4.62593 |
| periphFreq | Gen1 | male | 0.74946 | 0.10198 |
| totalDist | Gen1 | male | 0.42314 | 0.64157 |
| bw_1w | Gen1 | female | 0.00001 | 19.96487 |
| bw_4w | Gen1 | female | 0.00007 | 15.70274 |
| bw_7w | Gen1 | female | 0.00522 | 7.80164 |
| bw_sl | Gen1 | female | 0.00311 | 8.74096 |
| adj_heartW | Gen1 | female | 0.08176 | 3.02952 |
| adj_livrW | Gen1 | female | 0.21855 | 1.51386 |
| adj_adipW | Gen1 | female | 0.16883 | 1.89332 |
| centreDuration | Gen1 | female | 0.79135 | 0.06999 |
| periphFreq | Gen1 | female | 0.22805 | 1.453 |
| totalDist | Gen1 | female | 0.7718 | 0.08411 |
| LR | Gen1 | female | 0.96137 | 0.00235 |
| adj_EW | Gen1 | female | 0.60537 | 0.26697 |
| precocity | Gen1 | female | 0.00792 | 7.05164 |
| bw_1w | Gen2 | male | 0.00022 | 13.63858 |
| bw_4w | Gen2 | male | 0 | 38.18889 |
| bw_7w | Gen2 | male | 0 | 24.14779 |
| bw_sl | Gen2 | male | 0.00001 | 19.06475 |
| adj_heartW | Gen2 | male | 0.00868 | 6.88696 |

| | | | | |
|---|---|---|---|---|
| adj_livrW | Gen2 | male | 0.07319 | 3.21006 |
| adj_adipW | Gen2 | male | 0.2646 | 1.24451 |
| adj_testisW | Gen2 | male | 0.18355 | 1.76867 |
| centreDuration | Gen2 | male | 0.4461 | 0.58055 |
| periphFreq | Gen2 | male | 0.0241 | 5.08731 |
| totalDist | Gen2 | male | 0.65145 | 0.20408 |
| bw_1w | Gen2 | female | 0 | 25.20151 |
| bw_4w | Gen2 | female | 0 | 41.65775 |
| bw_7w | Gen2 | female | 0 | 31.61942 |
| bw_sl | Gen2 | female | 0 | 29.34427 |
| adj_heartW | Gen2 | female | 0.55856 | 0.34221 |
| adj_livrW | Gen2 | female | 0.30237 | 1.06371 |
| adj_adipW | Gen2 | female | 0.10166 | 2.67932 |
| centreDuration | Gen2 | female | 0.33818 | 0.91732 |
| periphFreq | Gen2 | female | 0.60951 | 0.26089 |
| totalDist | Gen2 | female | 0.97526 | 0.00096 |
| LR | Gen2 | female | 0.0328 | 4.55607 |
| adj_EW | Gen2 | female | 0.83055 | 0.0458 |
| precocity | Gen2 | female | 0.89131 | 0.01867 |

*Comment on additional file 6:*

When transformations were applied (purple distributions in **B.**, **C.**, **D.**) traits exhibited a more expected Gaussian distribution and the linear model used to estimate phenotypic variance components was therefore fitted on those normalized traits. Bodyweight at slaughter (bw_sl) and adjusted liver weight (adj_livrW) display a bimodal distribution in all generations, even after applying transformations. This is due to the strong sexual dimorphism that quails portray at adulthood. Performing Kruskal-Wallis tests in a sex specific way allowed taking this effect into consideration. Generally, laying rate (LR) and precocity were traits exhibiting the most deviations from normality. Shapiro-Wilk normality tests confirm that a variety of traits deviate from normality. Kruskal-Wallis tests evidence mean differences between epilines for BW from G1 onwards in females, and in G2 males.
