## Supplementary material 7 for "Transmission of an environmental modification in quails across three generations: changes in the contribution to phenotypic variability"

**Additional file 7:** Contribution of parameters to phenotypic variability per generation. **A.** Summary table of the linear model fitted for each trait with the best set of parameters kept after the stepAIC. The "trans" suffix was added when the trait was transformed during the normalization step. bw_1w: bodyweight at one week, bw_4w: bodyweight at four weeks, bw_7w: bodyweight at seven weeks, bw_sl: bodyweight at slaughter, adj_heartW: adjusted heart weight, adj_livrW: adjusted liver weight, adj_adipW: adjusted abdominal fat weight, LR: laying rate, adj_EW: adjusted egg weight, totalDist: total distance, centreDuration: duration at centre, periphFreq: frequency at periphery. **B.** Contribution of parameters (sex, family, epiline, reproductive status and their interactions) to phenotypic variability (in percentage) per generation. Interaction parameters were divided into two separate parameters: the interaction involving the epiline as one of the factors (interaction epi) and the interaction involving other parameters than the epiline (interaction non epi). **C.** Summary table of the estimates of the contribution of parameters (and standard deviation) to phenotypic variability (expressed in percentage). **D**. Model robustness ($R^2$) and correlations after cross validations. The coefficient of determination $R^2$ (blue) and the correlation between observed and predicted values in the Test set (yellow) were determined in 100 cross validations, to evaluate the predictive performance of linear models. The results show the distribution of $R^2$ and correlation values in each of the 100 cross validations, and are displayed per generation (x-axis) for each phenotype.

# A.

| Generation | Trait | Linear model equation |
|---|---|---|
| G0 | bw_1w | trait ~ sex + family |
| G1 | bw_1w | trait ~ sex + epiline + family + sex:epiline |
| G2 | bw_1w | trait ~ sex + epiline + family + epiline:family |
| G0 | bw_4w | trait ~ sex + epiline + family + epiline:family |
| G1 | bw_4w | trait ~ sex + epiline + family |
| G2 | bw_4w | trait ~ sex + epiline + family + epiline:family |
| G0 | bw_7w_trans | trait ~ sex + epiline + family + epiline:family |
| G1 | bw_7w_trans | trait ~ sex + epiline + family + epiline:family |
| G2 | bw_7w_trans | trait ~ sex + epiline + family + epiline:family |
| G0 | bw_sl_trans | trait ~ sex + epiline + family + epiline:family + sex:epiline |
| G1 | bw_sl_trans | trait ~ sex + epiline + family + epiline:family |
| G2 | bw_sl_trans | trait ~ sex + epiline + family + epiline:family |
| G0 | adj_heartW_trans | trait ~ sex + epiline + repro + sex:epiline + sex:repro |
| G1 | adj_heartW_trans | trait ~ sex + epiline + family + repro + epiline:family + epiline:repro |
| G2 | adj_heartW_trans | trait ~ sex + epiline + family + sex:epiline |
| G0 | adj_livrW_trans | trait ~ sex + family + repro + sex:repro |

| | | |
|---|---|---|
| G1 | adj_livrW_trans | trait ~ sex + repro + sex:repro |
| G2 | adj_livrW_trans | trait ~ sex + epiline + sex:epiline |
| G0 | adj_adipW_trans | trait ~ sex + epiline + family + repro + sex:epiline + epiline:repro + sex:repro |
| G1 | adj_adipW_trans | trait ~ sex + epiline + family + repro + epiline:family + epiline:repro + sex:repro |
| G2 | adj_adipW_trans | trait ~ sex + epiline + family + repro + epiline:family + epiline:repro |
| G1 | adj_testisW | trait ~ epiline + repro + epiline:repro |
| G2 | adj_testisW | trait ~ epiline + family + epiline:family |
| G0 | LR | trait ~ repro |
| G1 | LR | trait ~ family + repro |
| G2 | LR | trait ~ family |
| G0 | precocity_trans | trait ~ epiline + family + epiline:family |
| G1 | precocity_trans | trait ~ epiline |
| G2 | precocity_trans | trait ~ epiline + family |
| G0 | adj_EW_trans | trait ~ epiline + family + repro + epiline:family + epiline:repro |
| G1 | adj_EW_trans | trait ~ epiline + repro + epiline:repro |
| G2 | adj_EW_trans | trait ~ epiline + family + epiline:family |
| G0 | totalDist_trans | trait ~ sex + family + repro + sex:repro |
| G1 | totalDist_trans | trait ~ sex + epiline + family + repro + sex:epiline + sex:repro |
| G2 | totalDist_trans | trait ~ sex + epiline + family + epiline:family |
| G0 | centreDuration_trans | trait ~ sex + epiline + repro + sex:repro |
| G1 | centreDuration_trans | trait ~ sex + epiline + repro + sex:epiline + epiline:repro |
| G2 | centreDuration_trans | trait ~ repro |
| G0 | periphFreq_trans | trait ~ sex + family + repro + sex:repro |
| G1 | periphFreq_trans | trait ~ sex + epiline + family |
| G2 | periphFreq_trans | trait ~ sex + epiline |

B.

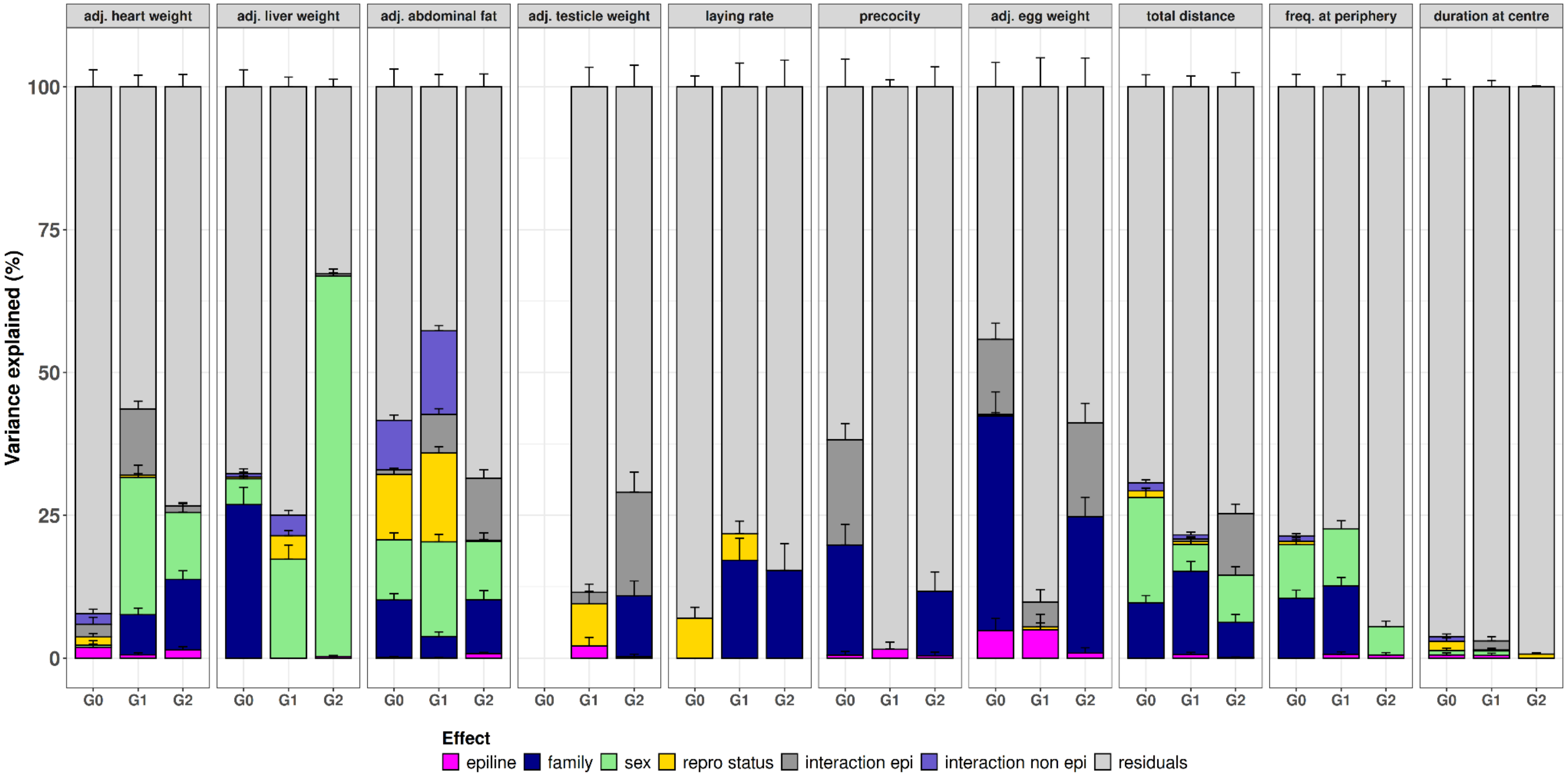
adj. heart weight
adj. liver weight
adj. abdominal fat
adj. testicle weight
laying rate
precocity
adj. egg weight
total distance
freq. at periphery
duration at centre
Variance explained (%)
100
75
50
25
0
G0 G1 G2
Effect
epiline
family
sex
repro status
interaction epi
interaction non epi
residuals

# C.

| Trait | gen | epiline | family | sex | repro | interaction epi | interaction non epi | residuals |
|---|---|---|---|---|---|---|---|---|
| adj. abdo fat | G0 | 0.14 (0.14) | 10.04 (1.13) | 10.54 (1 .22) | 11.43 (1.11) | 0.8 (0.33) | 8.69 (0.93) | 58.37 (3.12) |
| adj. abdo fat | G1 | 0.05 (0.07) | 3.75 (0.8) | 16.57 (1.26) | 15.53 (1.08) | 6.75 (1.01) | 14.64 (0.92) | 42.71 (2.17) |
| adj. abdo fat | G2 | 0.79 (0.21) | 9.43 (1.61) | 10.22 (1.47) | 0.21 (0.13) | 10.86 (1.49) | 0 (0) | 68.5 (2.24) |
| adj. heart weight | G0 | 1.91 (1.13) | 0 (0) | 0.34 (0.22) | 1.45 (0.61) | 2.25 (1.21) | 1.84 (0.75) | 92.21 (2.98) |
| adj. heart weight | G1 | 0.58 (0.36) | 7.03 (1.15) | 24.05 (2.12) | 0.41 (0.26) | 11.53 (1.4) | 0 (0) | 56.41 (2.05) |
| adj. heart weight | G2 | 1.42 (0.59) | 12.36 (1.51) | 11.72 (1.54) | 0 (0) | 1.18 (0.56) | 0 (0) | 73.32 (2.14) |
| adj. liver weight | G0 | 0 (0) | 26.87 (3.04) | 4.55 (1.73) | 0.28 (0.19) | 0 (0) | 0.57 (0.31) | 67.74 (2.95) |
| adj. liver weight | G1 | 0 (0) | 0 (0) | 17.3 (2.49) | 4.14 (0.88) | 0 (0) | 3.59 (0.83) | 74.96 (1.7) |
| adj. liver weight | G2 | 0.3 (0.17) | 0 (0) | 66.57 (1.23) | 0 (0) | 0.39 (0.19) | 0 (0) | 32.74 (1.32) |
| adj. testicle weight | G1 | 2.15 (1.48) | 0 (0) | - | 7.34 (2.19) | 2.01 (1.45) | 0 (0) | 88.49 (3.34) |
| adj. testicle weight | G2 | 0.27 (0.43) | 10.67 (2.57) | - | 0 (0) | 18.08 (3.53) | 0 (0) | 70.99 (3.77) |
| LR | G0 | 0 (0) | 0 (0) | - | 7.02 (1.89) | 0 (0) | 0 (0) | 92.98 (1.89) |
| LR | G1 | 0 (0) | 17.13 (3.86) | - | 4.63 (2.24) | 0 (0) | 0 (0) | 78.24 (4.14) |
| LR | G2 | 0 (0) | 15.38 (4.67) | - | 0 (0) | 0 (0) | 0 (0) | 84.62 (4.67) |
| adj. EW | G0 | 4.79 (2.15) | 37.6 (4.22) | - | 0.28 (0.31) | 13.13 (2.83) | 0 (0) | 44.2 (4.26) |
| adj. EW | G1 | 4.95 (2.73) | 0 (0) | - | 0.5 (0.68) | 4.36 (2.18) | 0 (0) | 90.18 (5.07) |
| adj. EW | G2 | 0.87 (0.94) | 23.91 (3.39) | - | 0 (0) | 16.44 (3.38) | 0 (0) | 58.79 (5.05) |
| precocity | G0 | 0.5 (0.66) | 19.31 (3.63) | - | 0 (0) | 18.41 (2.84) | 0 (0) | 61.78 (4.82) |

| | | | | | | | | |
|---|---|---|---|---|---|---|---|---|
| precocity | G1 | 1.54<br>(1.25) | 0<br>(0) | - | 0<br>(0) | 0<br>(0) | 0<br>(0) | 98.46<br>(1.25) |
| precocity | G2 | 0.42<br>(0.61) | 11.28<br>(3.39) | - | 0<br>(0) | 0<br>(0) | 0<br>(0) | 88.3<br>(3.48) |
| duration at centre | G0 | 0.57<br>(0.34) | 0<br>(0) | 0.76<br>(0.43) | 1.61<br>(0.59) | 0<br>(0) | 0.84<br>(0.44) | 96.22<br>(1.31) |
| duration at centre | G1 | 0.47<br>(0.38) | 0<br>(0) | 0.78<br>(0.47) | 0.17<br>(0.2) | 1.59<br>(0.73) | 0<br>(0) | 96.99<br>(1.12) |
| duration at centre | G2 | 0<br>(0) | 0<br>(0) | 0<br>(0) | 0.72<br>(0.21) | 0<br>(0) | 0<br>(0) | 99.28<br>(0.21) |
| freq. at periphery | G0 | 0<br>(0) | 10.45<br>(1.45) | 9.41<br>(1.23) | 0.57<br>(0.33) | 0<br>(0) | 0.92<br>(0.43) | 78.65<br>(2.2) |
| freq. at periphery | G1 | 0.68<br>(0.43) | 11.98<br>(1.46) | 9.99<br>(1.4) | 0<br>(0) | 0<br>(0) | 0<br>(0) | 77.35<br>(2.12) |
| freq. at periphery | G2 | 0.57<br>(0.4) | 0<br>(0) | 4.98<br>(0.95) | 0<br>(0) | 0<br>(0) | 0<br>(0) | 94.46<br>(1.02) |
| total distance | G0 | 0<br>(0) | 9.68<br>(1.27) | 18.46<br>(1.66) | 1.13<br>(0.46) | 0<br>(0) | 1.43<br>(0.51) | 69.3<br>(2.11) |
| total distance | G1 | 0.61<br>(0.39) | 14.55<br>(1.74) | 4.77<br>(1.14) | 0.48<br>(0.43) | 0.46<br>(0.35) | 0.71<br>(0.5) | 78.42<br>(1.88) |
| total distance | G2 | 0.07<br>(0.12) | 6.18<br>(1.38) | 8.29<br>(1.45) | 0<br>(0) | 10.74<br>(1.67) | 0<br>(0) | 74.71<br>(2.5) |

**D.**

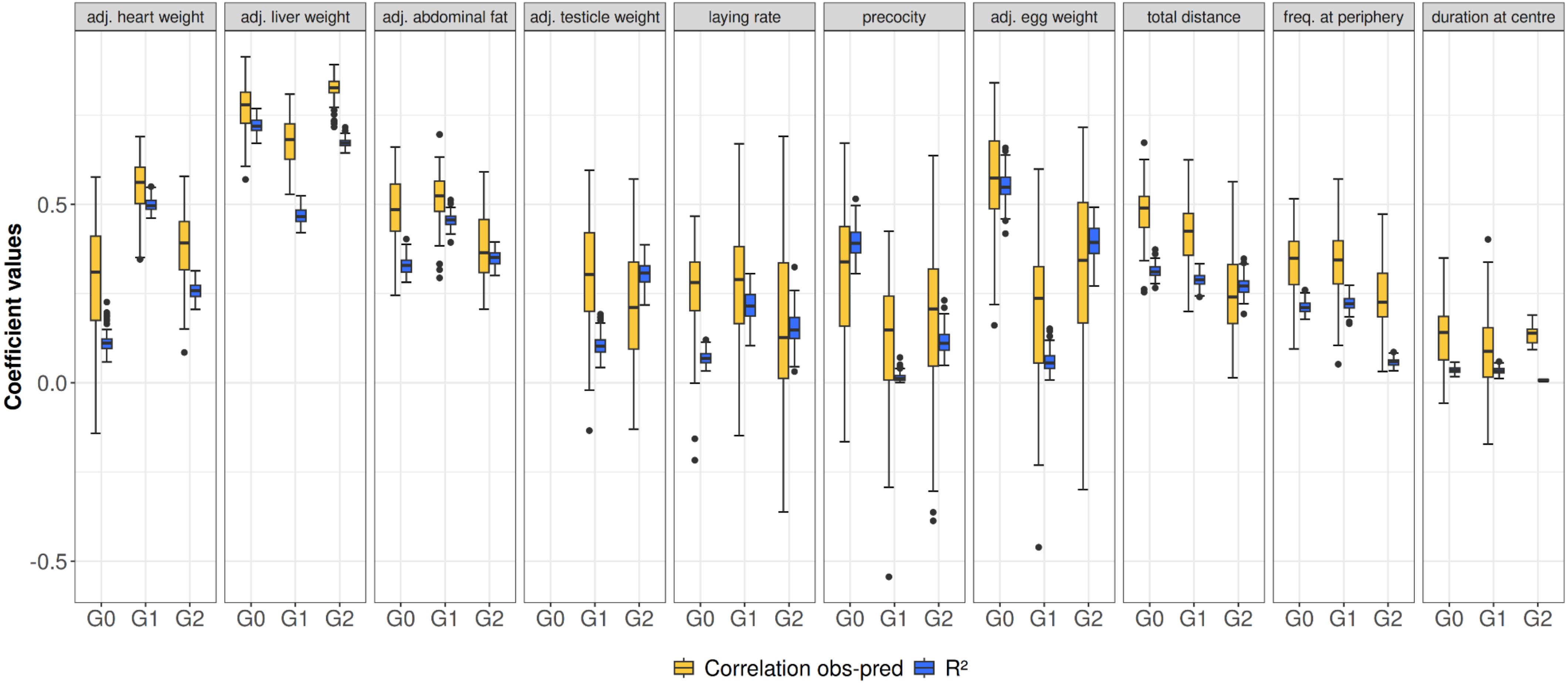

adj. heart weight
adj. liver weight
adj. abdominal fat
adj. testicle weight
laying rate
precocity
adj. egg weight
total distance
freq. at periphery
duration at centre
Coefficient values
0.5
0.0
-0.5
G0
G1
G2
Correlation obs-pred
R²

*Comment on additional file 7:*

Model fitting revealed that the epiline was a significant factor to keep for explaining a portion of phenotypic variability for several traits. In adjusted tissue weights like heart, abdominal fat and testicle, this parameter was kept after the stepAIC in all generations. For the liver, it was only found significant in G2. For production traits, the epiline was found significant in precocity and in adjusted egg weight at every generation but not for the laying rate. Finally, in behavioural traits, the epiline factor was kept in the models for total distance and frequency at periphery in G1 and G2 only, whereas it was kept in G0 and G1 for duration at centre. Although the epiline appears as a significant factor in the aforementioned traits, the estimates of the portion of variability explained remain below 5% in all, and negligible in most (inferior to 1% in adjusted abdominal fat, adjusted liver weight, total distance, frequency at periphery and duration at centre).

Besides, for all the traits presented in this figure, a substantial part of phenotypic variance remains uncaptured by our model, highlighting a strong limit in explanatory power and a limited confidence towards the estimates of the portion of phenotypic variability explained by the epiline. This is underlined by the large standard deviations found for all estimates, as well as the limited correlation and determination coefficients. The interaction parameter that includes the epiline shows higher estimates according to the trait and generation, reaching up to 16.44% in adjusted egg weight in G2, or 18.41 in precocity in G0 for example, yet the model for these two traits fails to explain half of the overall phenotypic variability (respectively 41.21% and 38.22%).
