## Supplementary material 8 for "Transmission of an environmental modification in quails across three generations: changes in the contribution to phenotypic variability"

**Additional file 8:** Simulations of a phenotype resembling body weight at slaughter over 5 generations. The trait was simulated as a polygenic trait (1000 QTL for 10000 chromosome segments) of mean 200, to resemble body weight at slaughter, variance 50 and an environmental variance of heritability 0.5 (as described in Karami et al. 2025). An initial founder population (PM) was simulated, it comprised 40 individuals, 20 males and 20 females, crossed with each other and each producing 10 progenies. These crosses initiated 20 founder families, as in our experimental design. From their offspring, 20 males and 40 females were selected, each male was mated with two females, therefore initiating the epilines (EPIP and EPIM). The males and females to mate were chosen to be from different founder families. Thereafter, mating was performed within each epiline independently and in mirror with the aim to cross different families in every generation. The simulation framework was repeated 100 times independently. The simulations were made using AlphaSimR (Faux et al. 2016, Gaynor et al. 2021).
**A.** Violin plot showing the distribution of the differences (in absolute) in phenotypic values between the two epilines, for each sex. The black dots are the 100 simulated values, the red dots are the values obtained from real data.

*Comment on additional file 8:*
Using simulation, we observe differences between epilines in phenotypic values for a number of simulations and at all generations. This result supports the conclusion made by David & Ricard 2023, that the difference in phenotypes can happen due to drift only. However, the differences in body weight at slaughter between epilines obtained with our simulations are lower than the ones observed with the real data, for generations 1 and 2 in females, and generation 2 in males, which concurs with our conclusions on the potential epiline effect on this trait.

**A.**

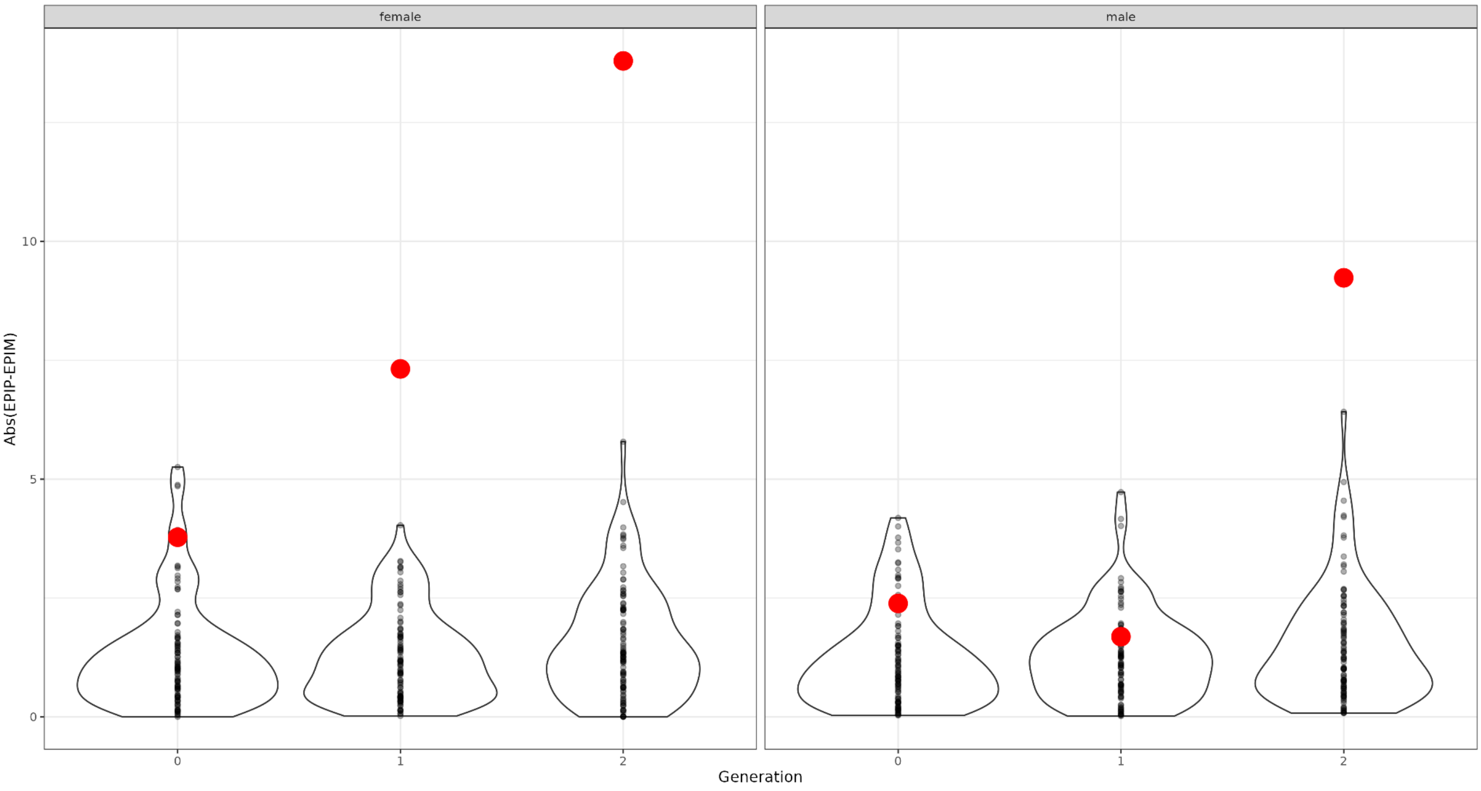
